# Distributed information encoding and decoding using self-organized spatial patterns

**DOI:** 10.1101/2022.06.04.494770

**Authors:** Jia Lu, Ryan Tsoi, Nan Luo, Yuanchi Ha, Shangying Wang, Minjun Kwak, Yasa Baig, Nicole Moiseyev, Shari Tian, Alison Zhang, Neil Zhenqiang Gong, Lingchong You

## Abstract

Dynamical systems often generate distinct outputs according to different initial conditions, and one can infer the corresponding input configuration given an output. This property captures the essence of information encoding and decoding. Here, we demonstrate the use of self-organized patterns, combined with machine learning, to achieve distributed information encoding and decoding. Our approach exploits a critical property of many natural pattern-formation systems: in repeated realizations, each initial configuration generates similar but not identical output patterns due to randomness in the patterning process. However, for sufficiently small randomness, different groups of patterns that arise from different initial configurations can be distinguished from one another. Modulating the pattern generation and machine learning model training can tune the tradeoff between encoding capacity and security. We further show that this strategy is applicable to non-biological dynamical systems and scalable by implementing the encoding and decoding of all characters of the standard English keyboard.

**Significance Statement:** Self-organized patterns are ubiquitous in biology. They arise from interactions in and between cells, and with the environment. These patterns are often used as a composite phenotype to distinguish cell states and environment conditions. Conceptually, pattern generation under an initial condition is encoding; discerning the initial condition from the pattern represents decoding. Inspired by these examples, we develop a scheme, integrating mathematical modeling and machine learning, to use self-organization for secure and accurate information encoding and decoding. We show that this strategy is applicable to non-biological dynamical systems. We further demonstrate the scalability of the scheme by generating a complete mapping of the standard English keyboard, allowing encoding of English text. Our work serves as an example of nature-inspired computation.

## Introduction

Information encoding is a process of converting information, such as text and images, from its original representation to an output format following defined rules. Dynamical systems have this information encoding capability as they can generate specific outputs according to given inputs. Conversely, decoding can be achieved if one can infer the input corresponding to an output. Depending on the system, decoding could be obvious, challenging, or impossible.

One example is to use cellular automaton (CA) that converts a grid of cells from a simple initial configuration into a self-organized sequence or spatial pattern according to a set of update rules(1). Wolfram proposed to use a chaotic rule to generate random sequences to encode information(2, 3). Here, the encoding is deterministic --each initial configuration corresponds to a unique output pattern. Because of the chaotic nature of the rule, however, decoding the input from a given output pattern is computationally prohibitive without prior knowledge of the update rules. As such, the system in theory can serve as the foundation for digital cryptography (4-8).

While making the encoding secure, however, the chaotic nature of the above example can limit its application. Like other dynamical systems exhibiting deterministic chaos, the final patterns generated by CA are extremely sensitive to perturbations and lack statistical regularities(9, 10). As such, a minute change in the initial configuration or the encoding process can lead to drastically different final patterns (a phenomenon termed the *avalanche effect*(11)). Unless the encoding and transmission are noise-free, the decoding is prone to errors *even if the rules are known* (12).

In contrast to these chaotic systems, many natural systems are convergent. That is, for the same or similar input configurations and environmental conditions, the final patterns share global similarity despite local variances. This property is sometimes referred to as “edge of chaos”(13). Examples are chemical reaction(14) and cortical networks(15). Many biological patterning systems also fall into this category. Despite minute variances, coat patterns are largely determined by animal genomes and allow identification of different species. In microbes, the same bacterial strain can grow into colonies with distinct shapes and sizes under different growth conditions(16, 17). Consequently, colony morphology can serve as a crude signature to distinguish environmental conditions and chemical cues, as well as the stage of infectious diseases (18, 19). Despite these empirical examples, the potential and limitations of information encoding and decoding using biological self-organization remain unexplored. Here, we use these systems to establish distributed information encoding. Coupled with machine learning (ML) mediated decoding, our system illustrates a scalable strategy for information encoding and decoding with quantifiable reliability and security (Figure 1A).

**Figure 1.**
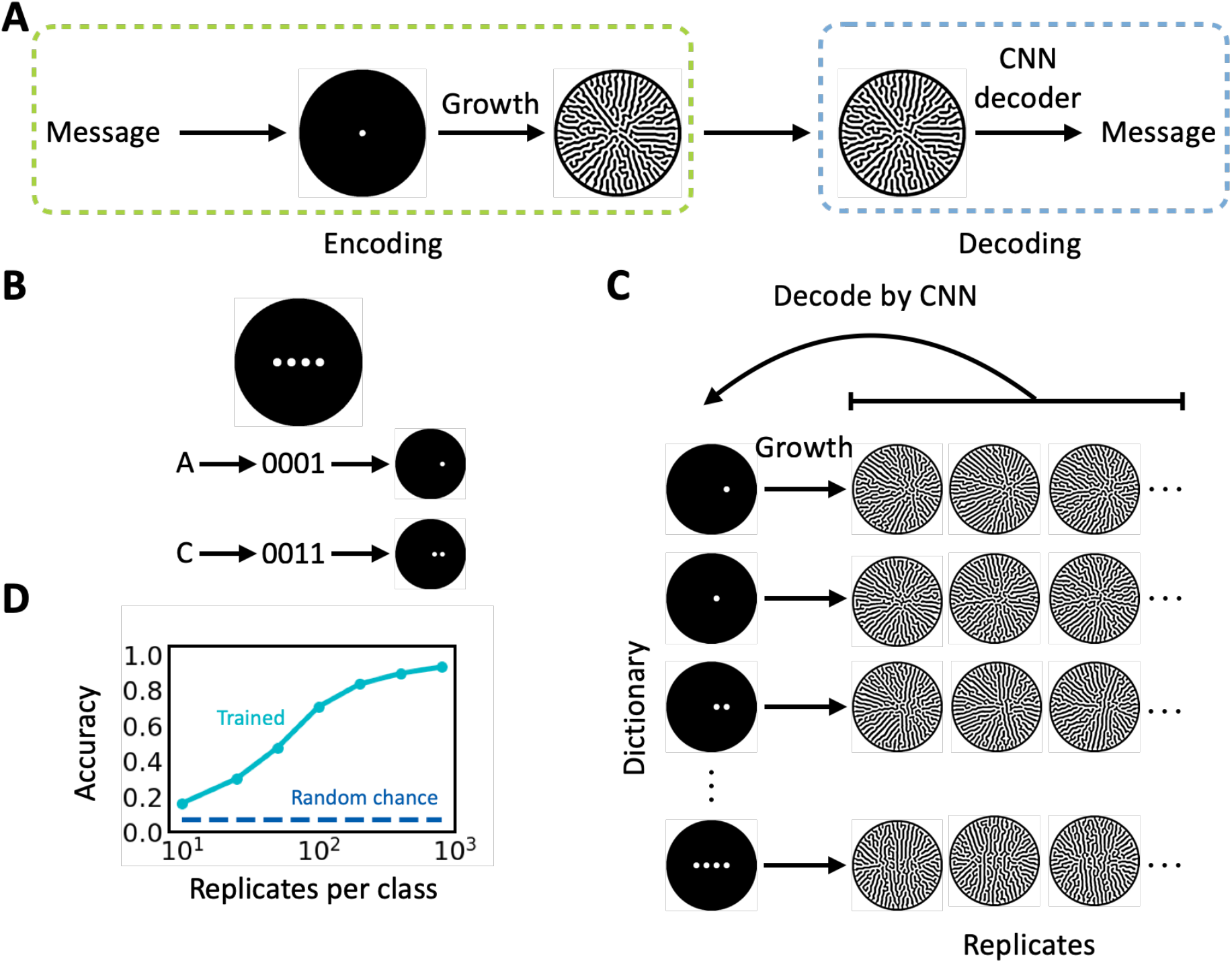
Distributed encoding and decoding using self-organized patterns. A. The encoding and decoding scheme. To encode, a message is converted into cell seeding configuration followed by colony growth, during which a colony pattern develops. To decode, the colony pattern of interest is fed into a trained CNN that converts the pattern into the original message. B. Predefined braille-like cell seeding arrangement. For a dictionary consisting of 15 characters (A-E and 0-9), we need a minimum 4-digit spot array (top). The characters (ex. “A” and “C”) are first converted into a 4-digit binary number, then converted into a seeding configuration. For a given digit, if it is 1, cells are “inoculated” within the corresponding spot and if it is a 0, no cell is inoculated. C. One-to-many mapping between seeding configuration and spatial patterns. Pattern formation is subject to minor biological noise, which includes heterogeneity in cell seeding, external perturbation and variability in cell phenotype during growth process. The noise is amplified by the branching mechanism. Hence patterns evolved from the same configuration share qualitative similarity but are different in detail. A well-trained CNN should navigate through this mapping and be able to decode the patterns as the corresponding character. For CNN training, the dataset is composed of equal number of replicates of patterns developed from all seeding configurations. D. Relationship between the number of replicates of the training set and CNN accuracy. The CNN was trained on a balanced dataset that contains 15 distinct characters. The numerical simulation used the default parameter values (see “Mathematical modeling” in Methods) and intermediate growth noise (signal-to-noise ratio = 3.5). The CNN decoding accuracy increases as the number of available replicates increases. The accuracy is significantly higher than random chance (1 / the size of the dictionary).

## Results

### Criteria for Choosing an Encoding System

Any dynamical systems, including those generating self-organized patterns, can serve as the foundation for information encoding and decoding. However, to ensure secure encoding and reliable decoding, we reason that the system dynamics need to meet a set of heuristic criteria. First, the output patterns are sufficiently complex and diverse such that different initial configurations would generate distinguishable output patterns. Second, the pattern generation is subject to stochasticity but remains convergent. That is, in repeated pattern generation processes, the same initial configuration with small noise or perturbations should generate output patterns that are approximately the same but differ in minor details. Importantly, the differences between patterns generated from replicated simulations should be smaller than those between patterns generated from different inputs. Third, while different groups of patterns arising from different initial conditions can be decoded by a properly constructed decoder, their differences are difficult to discern by naked eyes. We note that the degree by which different groups of patterns can be distinguished often has to be established empirically (if a reliable decoder can indeed be constructed).

As a proof of principle, we focus on a coarse-grained model of self-organized pattern formation (Figure 1, also see “Mathematical modeling” in Methods). The model was developed to simulate qualitative aspects of branching dynamics of *Pseudomonas aeruginosa* colony growth (20). In it, each simulation initiates from a pre-defined cell seeding configuration and the cells develop into a branching colony (Figure S1). The patterning process is influenced by two sources of random noise. One comes from the variability in the initial distribution of seeding cells; the other comes from the underlying growth kinetics. With appropriate choice of parameters (including noise levels), the patterning dynamics satisfy all criteria listed above.

In addition, another rationale for choosing this model is its simplicity and versatility. It can generate diverse patterns by adjusting model parameters and be solved in a computationally efficient manner. These features allow us to probe this platform’s security, reliability, and scalability (see “Tradeoff among encoding capacity, security, and decoding reliability”).

### Distributed Encoding and Decoding by Spatial Patterns

To demonstrate encoding, we represent a dictionary of 15 characters — letters A-E and numbers 0-9 — using binary numbers 0001-1111 (Table S1). Each binary number then corresponds to a seeding configuration of cells in a braille-like array at time 0 (Figure 1B): a digit “1” corresponds to a spot seeding indicating the presence of cells, whereas a digit “0” indicates no cells. In each simulation, the colony grows from its initial configuration into a final pattern. As mentioned above, the simulation is subject to two noise sources: the variability in seeding and during growth. The former could originate from the marginal but unavoidable uneven cell seeding, and the latter could originate from inherent heterogeneity of cell gene expression, motility, or small external perturbation. Therefore, repeated simulations from the same initial seeding configuration generate similar final patterns with minor differences, which *collectively* encode the identity of the input configuration (Figure 1C). We chose to encode in seeding configuration because of its simplicity, one may also choose to encode in other parameters influencing pattern formation.

We configure our simulations such that neither the mapping between the initial configurations and the colony patterns nor the difference between patterns corresponding to different inputs is obvious to the naked eye. To allow reliable decoding, we need a robust method to navigate through this visual complexity. A direct method is brute-force search, whereby all the possible patterns for each initial configuration are simulated to establish an empirical mapping between the input and the output. While apparently straightforward, this approach is computationally prohibitive and impractical because the training patterns are 8-bit, 80 pixels × 80 pixels grayscale images, resulting in up to 2^8×80×80^ ∼ 10^15412^ possible patterns.

Alternatively, image classification using convolutional neural networks (CNNs) has been successful for numerous applications (21-23). Through observing sufficient examples, a CNN learns to cluster images by their categories. Here we built a CNN to decode the colony patterns via multiclass classification (Figure S2, see “CNN training” in Methods). During training, our CNN decoder takes pattern images (generated by repeated simulations) as input and updates its trainable parameters to classify patterns based on initial seeding configurations. With sufficient replicates in each class, our trained CNN was able to distinguish patterns corresponding to the 15 characters with high accuracy (Figure 1D). For instance, greater than 93% of decoding accuracy can be achieved by having 800 replicate patterns in the training set. We note that this decoding approach is data-driven; other algorithms such as decision tree (24) and support vector machines (25) may also be used.

In an actual application of this encoding/decoding strategy, we assume the channel is public while the pattern generator, model parameters, training set, and the trained CNNs are private to the end users (Figure 1A). The recipient chooses the correct, trained CNN to decode a pattern according to the model parameters transmitted through another private channel (not shown in the figure) as prior knowledge.

### Tradeoff Among Encoding Capacity, Security, and Decoding Reliability

In this platform, we aim to maximize the capability of the patterns to encode information, termed *encoding capacity*, and our platform’s robustness against data leakage to unauthorized parties, termed *encoding security*. We consider a system has higher encoding capacity if it can encode more characters correctly with adequate data, while we consider our encoding scheme being more secure when the attacker cannot build a successful decoder from the leaked data. For example, the accuracy of a separate decoder built on only 10 replicates per class drops to less than 20% (Figure 1D), which is only slightly better than random guessing (1/15). Note that the efficacy of our platform depends on the complexity of the generated patterns, our desired accuracy, and the amount of available training data.

We can tune our scheme’s performance by modulating parameters in the pattern generation model, i.e., the relative acting distance and magnitude of colony expansion versus repulsion processes. Large relative distance and magnitude (i.e., higher colony expansion) result in thick branches, whereas small relative distance and magnitude (i.e., higher repulsion) result in thin, sparse branches. In extreme cases, these conditions can result in large disks or small circular colonies, respectively. When these two forces are intermediate and comparable, the system generates branching colonies. We constructed 16 simulated training datasets of diverse patterns by tuning these two parameters (Figure S3A). Based on their final appearance, we categorized our results into three subgroups: disk-like, trivial (final pattern is identical to initial configuration), and branching. Disk-like colonies cannot be distinguished regardless of the training data size—thus, the input information was obscured and “lost” after growth (Figure S3B-D). Conversely, trivial patterns allow perfect but insecure decoding since the reverse mapping is obvious. Ultimately, the intricate branching patterns allow secure encoding and reliable decoding as demonstrated previously.

We can also modulate encoding capacity and security by tuning the noise during the patterning process. Without noise, one pattern per input is sufficient for perfect decoding as long as output patterns are distinguishable (Figure 2A). Too much noise would introduce too many variations in the replicate patterns generated from each input. If these intra-category variations (between replicate patterns) approach or exceed the inter-category differences (between sets of patterns corresponding to different inputs), the decoding accuracy would deteriorate significantly (Figure 2A). Depending on the magnitude of the noise, this loss in accuracy can be alleviated by increasing the number of replicate patterns per class. A similar tradeoff exists for other parameters as well, such as the spacing between spots in the initial configuration (Figure 2B). When spacing decreases, patterns grown from different configurations appear more alike and indistinguishable. Moreover, increasing dictionary size with all else being equal would also reduce the decoding accuracy (Figure 2C). Again, expanding the number of replicate patterns per class can compensate for losses in accuracy, thus increasing the encoding capacity (Figure 2D). Similar tradeoff was also observed in patterns arrested from growth at different time points (see “Temporal information encoding and decoding” in Supplementary Information).

**Figure 2.**
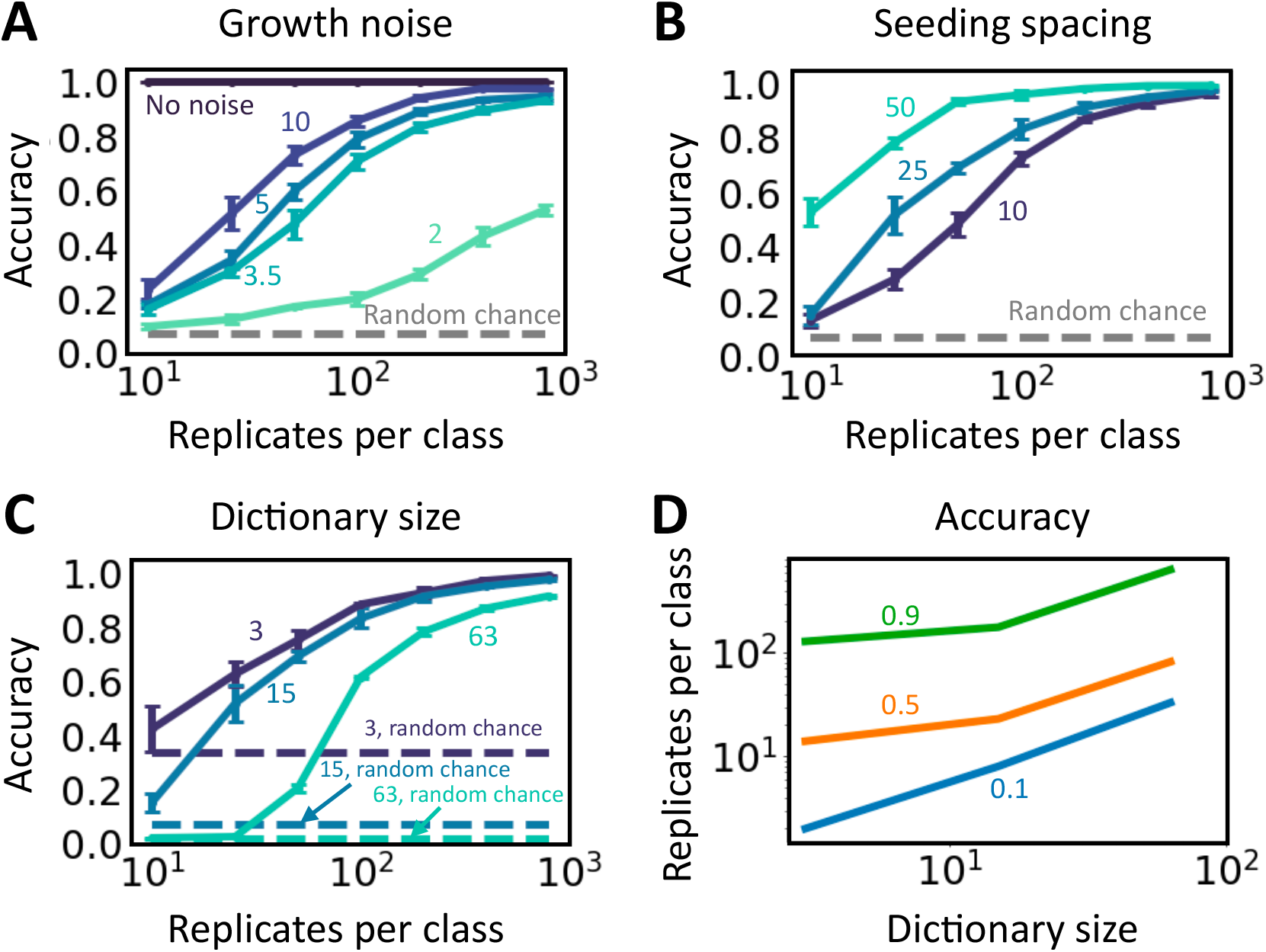
Tradeoff between encoding capacity, security, and decoding reliability. We present the prediction performance of CNNs trained on branching patterns with different parameterizations. Specifically, A. We fixed the seeding noise such that the only source of noise is growth. The magnitude of growth noise is modulated through changing the signal-to-noise ratio (SNR) of the growth kernel. The higher SNR is, the lower the noise level is. We present the results on datasets with no noise, SNR = 2, 3.5, 5 and 10 respectively. B. We simulated patterns using seeding spacing = 10, 25, and 50 respectively, which represent from small to large spacing. As the spacing increases, patterns corresponding to different initial configurations become more dissimilar. C. We simulated datasets of 3, 15, and 63 characters using 2-, 4- and 6-bit predefined braille-like seeding arrays respectively, while keeping all else as the default. Overall, the decoding accuracy increases as the number of replicates per class increases, and it significantly exceeds the corresponding accuracy by random guessing. The only exception is in the absence of growth noise, in which case the patterns are identical thus the decoding is trivial. Notably, when the patterns become more complicated (ex. larger growth noise, smaller spacing, or larger dictionary), more data are required to reach the same accuracy. D. Required training replicates per class as a function of dictionary size. The green, orange, and blue lines represent accuracy of 0.9, 0.5 and 0.1, respectively. The required data size increases exponentially as the desired accuracy increases.

In principle, the encoding-decoding scheme is applicable to any dynamical systems where the input-output mapping satisfies the criteria listed above. To illustrate this point, we chose an elementary cellular automaton model with weakly chaotic dynamics (9) (see “Encoding and Decoding using Elementary Cellular Automaton” in Supplemental Information). Given the set of rules, we chose the model parameters (including noise levels) such that the resulting dynamics can allow secure encoding and reliable decoding. Again, we encoded characters in binary numbers, which is then converted into 1D initial configuration in a similar manner as in 2D. Noise was imposed on the initial sequence, and the latter develops into a final sequence following the evolution rules (Figure S4). A feedforward neural network was trained to code the final sequence. As expected, higher complexity leads to worse decoding accuracy, and it can be remedied by increasing training data size (Figure S5).

### Enhancing Encoding Security and Integrity

To enhance security, we evaluated utilizing encryption to prevent unauthorized access during communication. A secret key is implemented during encoding and successful decoding requires the correct key (Figure 3A). For pattern formation systems, the geometry of the patterning domain is a feasible choice of secret key as it can influence the patterning process and is easily tunable (26-28). In our system, the boundary suppresses bacteria colonization, and the strength of the impact decreases exponentially as the distance from the focal niche to the boundary increases (see “Encryption dataset generation” in Methods). As such, the boundary exhibits a time-invariant, long-range, and weak inhibitive force on colony expansion. As this force is anisotropic due to asymmetric boundary geometry, the patterns are encrypted by the domain shape.

**Figure 3.**
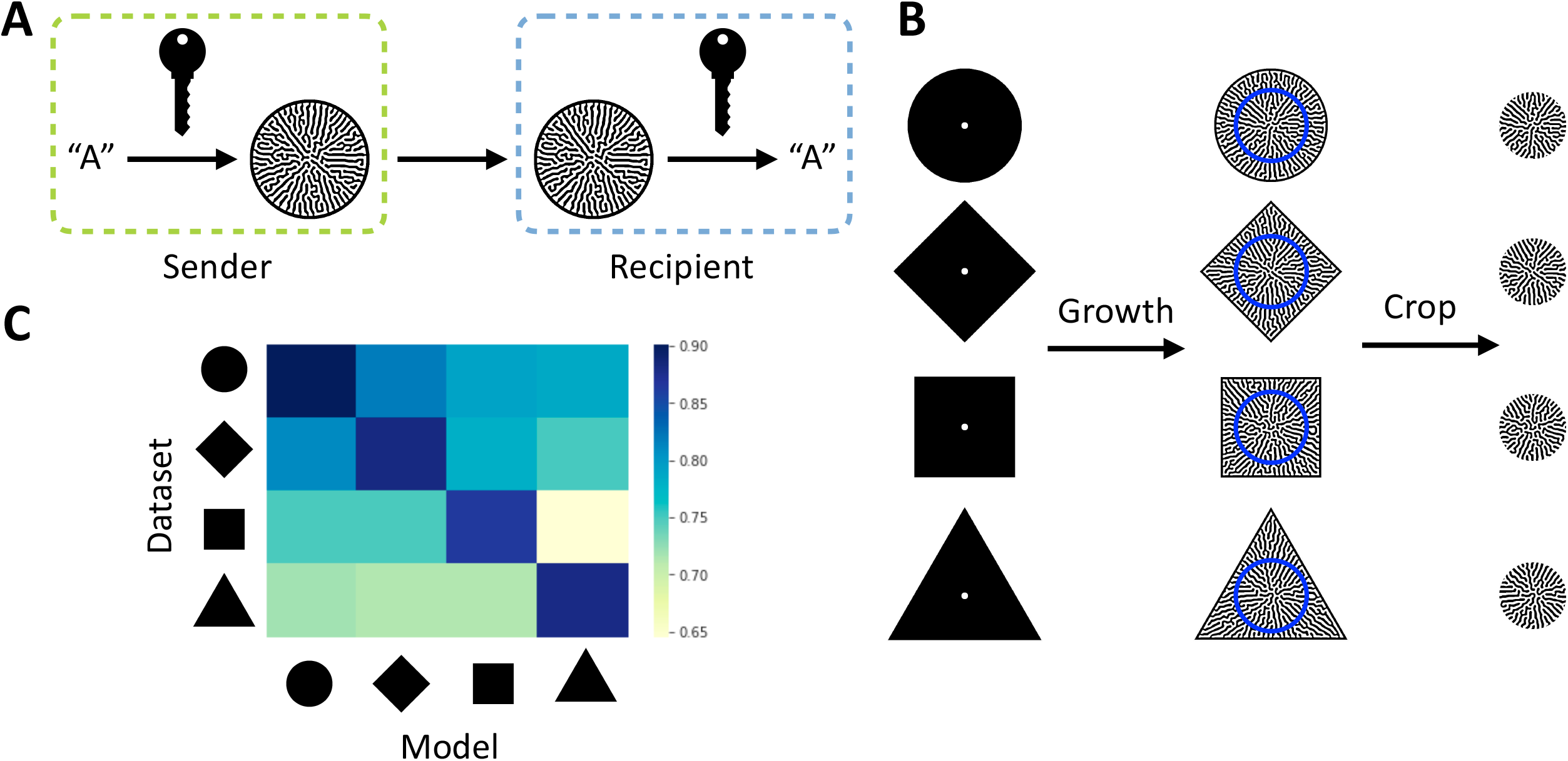
Encryption using growing domain shape as the secret key. A. Encryption scheme. A secret key is used to convert a message (ex. “A”) to a self-organized pattern, and the knowledge of it is required to reliably convert the pattern back to the original message. For our ML-mediated decoding method, the information on the secret key allows the designated recipient to choose the correct, trained CNN to decode the received pattern. B. Training data generation and preprocessing. For each encoding character, we computationally seeded cells on growing domains of different shapes (left) and let them grow into spatial patterns over the entire field (middle). The centers of the colonies (within the blue circles) were cropped to remove the information of the growth domain (right), and then used for CNN training. C. Effectiveness of encryption when growth domain shape is the secret key. Four CNN models were trained independently on datasets encrypted by circular, diamond, square and triangular growth domains respectively. The heatmap shows their decoding accuracies on each dataset. Only the model trained on the corresponding dataset can decode at the highest accuracy.

To test this notion, we generated patterns within different boundary shapes. For each shape, the resulting patterns would occupy the entire space. We removed the information of the boundary in the output by cropping out a smaller, circular area at the center of each pattern (Figure 3B). We found that only the decoders trained on the correct datasets can decode at high accuracy (Figure 3C), indicating that knowledge of the domain shape (i.e., the secret key) is critical for selecting the right CNN decoder to accurately decode. Similarly, we evaluated the potential of other secret key choices, such as the seeding spacing (Figure S7) and patterning domain size.

We have also considered the threat to information integrity during communication, in which the attackers could alter the output patterns or replace them with fake ones, thus deceiving the intended information receiver. We demonstrated that the noise in the patterning dynamics could be used to ensure the integrity (see “Authenticating patterns using noise signatures” in Supplementary Information). Briefly, the noise leaves a unique signature for each correct pattern, which can be used to authenticate a received pattern.

### Improving Decoding Performance by Ensemble Learning

All else being equal, the reliability of decoding can be improved by increasing the number of replicates per class when training the decoder. However, the degree of improvement diminishes for an increasing number of replicates (Figure 2C). For instance, for a dictionary size of 63, the decoding accuracy increases ∼30 folds by increasing the number of replicates from 10 to 100; it only increases ∼1.5 fold by increasing from 100 to 800. To more effectively use the available data, we adopted ensemble learning – a class of machine learning techniques (29-31).

Staked generalization combines the knowledge learned by individual, base ML models for better prediction (32-35). We first trained multiple base CNN decoders on a dataset with random initialization using the same protocol in the previous sections, then trained an ensemble decoder to combine their prediction capabilities. The ensemble model was then used for final decoding (Figure 4A, see “Ensemble learning and uncertainty estimation” in Methods). For patterns generated with moderate growth noise, the prediction performance of the ensemble decoder excels that of the base models for up to 22% in accuracy (Figure 4B). Receiver operating characteristic (ROC) curves and confusion matrices also show significant improvement with ensemble model (Figure 4C, Figure S8 - 9). As expected, the ensemble model generally outperforms the base ones when intermediate data are available but demonstrates marginal improvement with adequate or scarce data. This is expected because when intermediate data are available, the individual base models are diversified due to random initialization. However, when adequate data are available, each base model individually decodes with high accuracy, leaving little room for improvement. Conversely, when data are scarce, the base decoders barely learn such that integrating their results provides little insight. This final aspect implies encoding security against minor data leakage. Additionally, considerable improvement can be achieved with a simple logistic regression (LR) model, and more base models leads to better ensemble performance (Figure S10). In addition to stacking, we have also shown that majority voting can improve the decoding accuracy (Figure 4D, Figure S11). Multiple patterns corresponding to the same character were decoded using the same CNN, and the most voted prediction was used as the final prediction.

**Figure 4.**
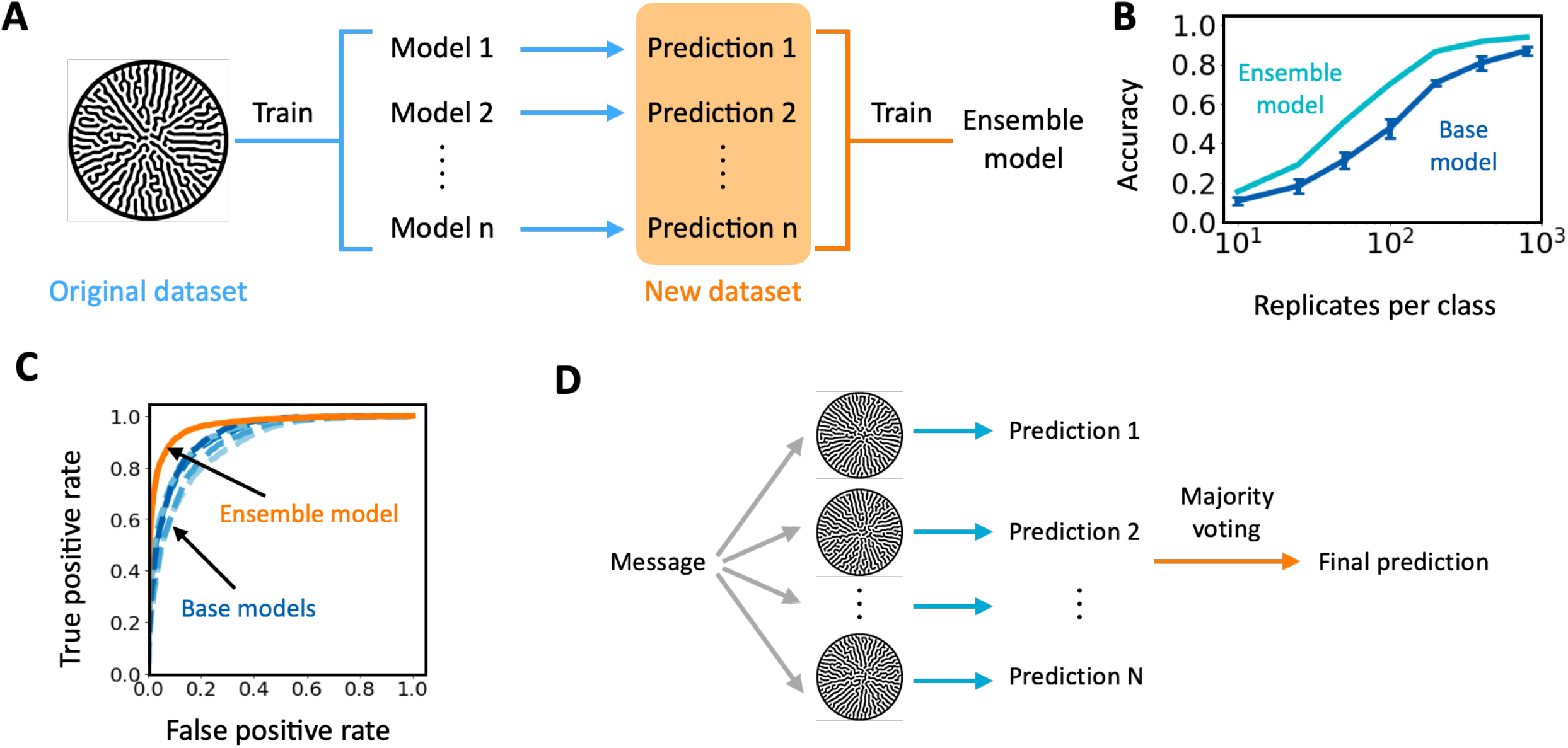
Using ensemble learning to improve decoding accuracy. A. Training procedure of the ensemble model. The training is done in two steps. First, we train multiple base CNN decoders on a dataset as described in the previous sections. Then their predictions on the training set and the corresponding class labels constitute a new dataset. In the second step, we train an ensemble model from stretch using the new dataset. B. Decoding accuracy of ensemble and base models. Here, a LR ensemble model was trained with five base models. The ensemble model outperforms the base models regardless of the training data size. Notable improvement in accuracy occurs when moderate amount of data was available for training, whereas the improvement is less significant with adequate or scarce data. C. ROC curve of ensemble and base models (orange: ensemble model; shades of blue: base models). The ROC curves were computed for each encoding character and then averaged over all classes to reflect the overall performance of the decoders. The area under the ROC curve (AUC ROC) of the ensemble model is 0.963. AUC ROC of the base models are 0.881, 0.893, 0.920, 0.925 and 0.924. The models were trained on a dataset with 100 replicated per class. D. Schematic of the majority voting algorithm. Instead of using only one pattern for communication, the sender would generate and send out multiple patterns representing the same message. Due to the randomness in the patterning process, these patterns appear similar but differ in detail. The recipient would use a trained decoder to decode each pattern and obtain the corresponding predictions. The most popular prediction will be used as the final prediction.

Ensemble learning not only improves the decoding accuracy, but also sheds light on the prediction uncertainty. According to Lakshminarayanan *et al*, the base models trained with random initialization explore the entirely different modes of function space(36), thus their independent predictions can be used to estimate well-calibrated uncertainty(37). We adopted this notion and estimated decoding uncertainty through multiple metrics, including log likelihood, mean squared error (MSE), top-1 and top-5 errors (see “Ensemble learning and uncertainty estimation” in Methods). Higher metric value indicates larger uncertainty or lower confidence. As expected, the uncertainty reduces as more training data are available (Figure S12). Having more base models does not necessarily reduce the uncertainty (Table S2).

The information on uncertainty can assist decision making on whether to accept the decoding result or not. Depending on the application, the end users could define a dichotomization accuracy and uncertainty. For a decoded character, if the decoding accuracy (a priori knowledge) and the uncertainty are inferior to the suggested threshold, the recipient should reject the pattern and request a new one. Otherwise, the decoded character can be accepted.

### Distributed Encoding of English in Emorfi

Our distributed encoding-decoding platform is scalable for practical applications. We constructed 100 sets of patterns to encode all printable ASCII characters including English letters in upper and lower cases, digits, punctuations, and whitespaces (Figure 5A, Appendix A). A 7-bit seeding array was used to create the training dataset, in which, 100 of the unique initial configurations corresponded to the printable characters. Each of the initial configurations was then used to generate 1000 patterns. When encoding text, each character is represented by one or multiple patterns that are then arranged to assemble a video (Figure 5B). We term this collection of patterns *Emorfi*, which represents a new, digitally generated coding scheme.

**Figure 5.**
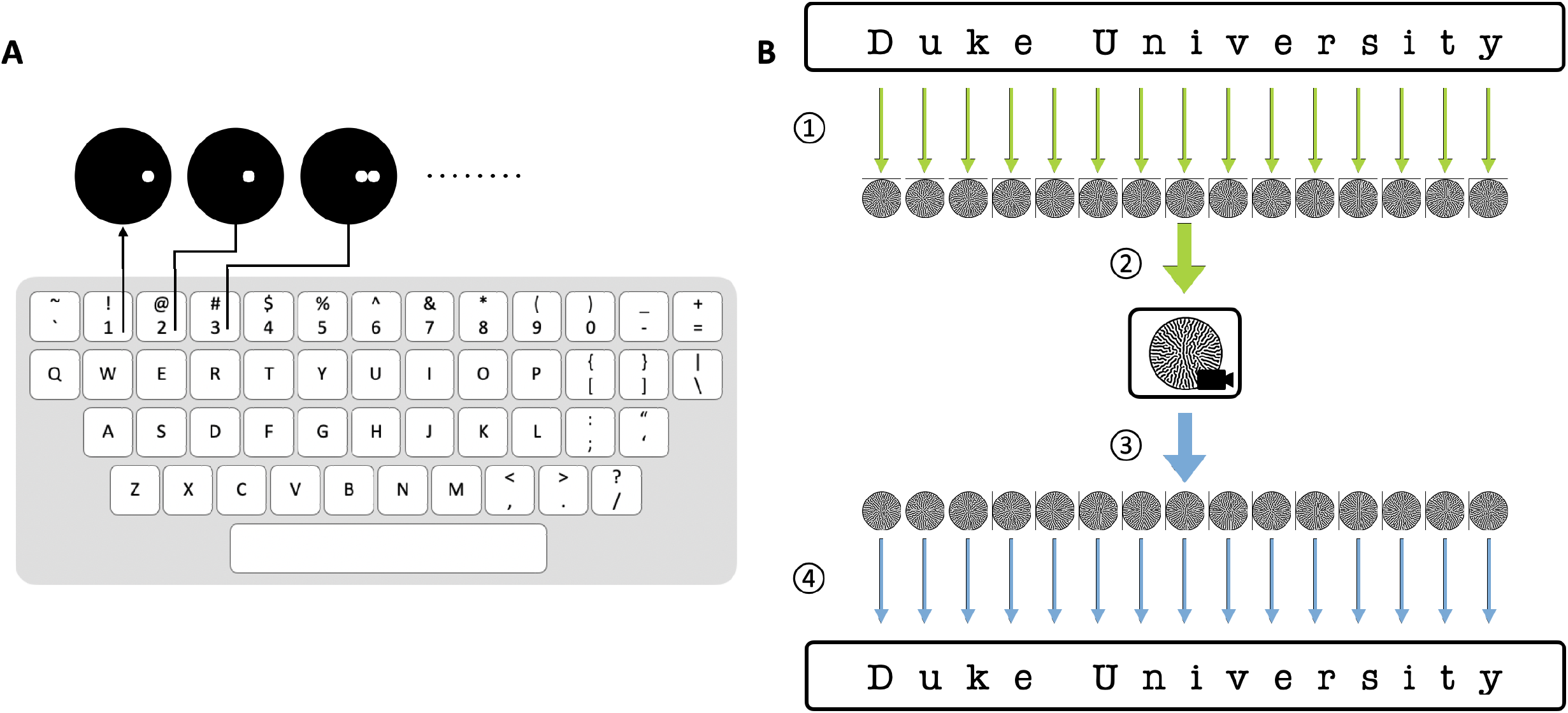
Encoding text in Emorfi A. Each of the 100 printable ASCII characters is represented by a unique initial configuration. 95 of them are shown on the keyboard, and 5 other printable whitespace characters (tab, linefeed, return, vertical tab, and formfeed) are not shown here. In the training set, each character maps to 1000 patterns. The collection of patterns, as well as subsequent ones to be generated, constitutes Emorfi. B. A piece of text could be encoded as a video and decoded using ensemble method. ⍰ Each character in the text is translated to a corresponding pattern. ⍰ The images are arranged in order and assembled into a video that can be used for commincation. ⍰ To decode, each frame is retreived from the video. ⍰ The patterns are decoded sequentially, representing the decoded text.

By doing so, all standard English text can be encoded in Emorfi and decoded back. For instance, we encoded the public speech “I have a dream” by Martin Luther King Jr. containing 8869 individual characters as a video (Movie S1). Accommodating majority voting, each character was represented by five different patterns. The video was decoded with 99.8% accuracy (Appendix B). The same approach was also used to encode the poem “Auguries of Innocence” by William Blake as video (Movie S2) and decoded with 99.6% accuracy (Appendix C). In another example, using a 5-bit seeding array, we encoded GFP protein sequence (238 amino acids) as a video (Movie S3) and decoded at 100% accuracy (Appendix D).

## Discussion

Our encoding and decoding framework is applicable to diverse dynamics systems, as long as they have three key properties: i) an approximately convergent mapping between initial input and output, ii) complex output signals, and iii) the output patterns are difficult to distinguish to the naked eye. While past studies have explored the possibility of using chaos to encode information and to provide security (38-40), unavoidable noise and error in numerical simulation (e.g., finite precision computing) or transmission (e.g., channel noise) can alter the output despite these systems being deterministic. In contrast, the convergent nature of our system ensures patterns that originate from the same initial configurations share common features (recognizable by a trained NN) despite small variances. Though noise is often considered undesirable in biological studies — such as masking ground truth (41-43) or disrupting interactions between components (44, 45)— we take advantage of the variance in our system to ensure information security and to authenticate each pattern.

These criteria together contribute to the sufficient encoding capacity and tunable information security of our platform. Many systems satisfy these criteria. With appropriate parameterization and boundary conditions, many reaction-diffusion models exhibit considerable robustness in output patterns and sensitivity to initial conditions (46, 47). In addition to the example we demonstrated (Figure 2 and S4-S5), many CA models with asynchrony update rules also show convergence (48, 49). Biological systems, such as biofilm morphology, butterfly wing scale pattern, and human fingerprint, have also evolved to exhibit common features but vary in detail. Their convergent nature results from the rich multiscale, multidimensional interactions between different system components, such as chemical reactions and diffusion, gene circuits, and cell-cell interactions (50-54).

However, our work does bring up a fundamental question: given a dynamical system with stochasticity, how do we know the dynamics are convergent enough while the output signals from different initial conditions are also distinguishable? We suspect that the question has to be addressed empirically for each specific system. In ours, each initial configuration generates an ensemble of output patterns following a distribution (visualized using t-SNE in Figure S13). It is difficult to determine this distribution by solely inspecting the pattern generation model, even if parameters and noise magnitudes are known. However, whether each distribution corresponding to an input can be distinguished from another distribution arising from another input is established by ML. In essence, the trained CNN provides an empirical estimate on the extent by which the pattern generation is convergent. To this end, our work has implications for quantifying the convergence for a dynamical system by using ML.

As we have demonstrated with Emorfi, the patter-based encoding-decoding platform is *scalable and generalizable* for information in various formats. We envision the platform could be extended to other languages, such as alphabetic languages with different letters or diacritics (e.g., French, Hebrew) and logographic scripts consisted of thousands of characters (e.g., Chinese, Japanese). It could also be applicable for communicating science and protecting intellectual properties by incorporating Greek alphabet, mathematical symbols, nucleic acid bases and etc.

## Methods

### Mathematical Modeling

The simple colony pattern generation model accounts for several driving forces. In particular, it uses a kernel-based method to capture the high level positive (expansion) and negative (inhibition) effects on patterning, regardless of the specific mechanism. The model is formulated as the following equations:

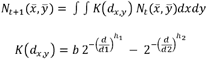

Here, N is the colonization of the bacteria over the growing medium, K is the growth kernel that is the addition of the expansion and (negative) repulsion kernels. b is the relative magnitude of expansion to repulsion, d_1_ and d_2_ are the distances that characterize half of the maximum effect of expansion and repulsion respectively. *d*_*x,y*_ is the distance of the focal point to 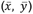. We used *d*_1_*/d*_2_ = 0.4, *h*_1_ = 1000, *h*_2_ = 2000, *b* = 6.5 as the default parameter values unless otherwise mentioned. This parameter set generates complex branching patterns.

To adapt the published model for our study, we made several modifications. First, we implemented various seeding configuration, such as the spot seeding arrangement for encoding binary representations of characters (Figure 1B). The size and spacing of the spots were subjected to modulation. As the default setting, we used spacing = 15 and spot radius = 5. Second, we implemented white Gaussian noise with varying signal-to-noise (SNR) ratios to the growth kernel at each time step. The noise mimics the heterogeneity and small perturbations in growth. We also implemented uneven cell seeding by assigning random intensities drawn from a truncated Gaussian distribution (mean = 0.5, deviation varies) to the pixels within the spot configurations. Both noise sources contribute to the variation in patterns given the same model parameters and initial configurations. As default, we used random seeding without growth noise.

The model was implemented in MATLAB 2017b and solved numerically. The simulation terminates once the colony stops growing. The simulation outputs an 8 bit, 451 pixels × 451 pixels greyscale image. Except the encryption experiments, the patterns were formed on a circular growth domain of a diameter of 451 pixels.

### CNN Training

For CNN training, we numerically simulated datasets with equal number of replicates for each encoding character. For evaluation, test datasets made of 100 replicates per class were used. The pattern images were rescaled to 80 pixels × 80 pixels before training or testing.

The CNN (Figure S2) and the ensemble model (Figure 4A) were implemented in Python 3, TensorFlow 1.15.2, and Keras 2.4.0. The CNN uses pattern images as inputs and outputs N features, where N is the dictionary size. It consists of two convolutions, each followed by max pooling and rectified linear unit (ReLU). Then their output is passed onto two fully connected layers, followed by ReLU and softmax respectively. Here the softmax function turns it into categorical probabilities. For training, we used Glorot normal initializer, categorical cross entropy loss and Adam optimization algorithm with learning rate subject to tuning. Keras early stopping function was also implemented to stop the training once the loss metric stopped improving. We carried out hyperparameter tuning (including learning rate, batch size, early stopping patience and delta) to obtain the best performing models for analysis. The data generation and training were conducted on Duke Compute Cluster and Google Cloud Platform.

### Encryption Dataset Generation

The geometry of the growth domain impacts the growth and pattern formation through exerting a negative effect on the colony in the vicinity of the boundary, such that the colony does not reach the edge. The plate influence is formulated as:

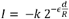

Here, *d* is the Euclidean distance of the focal niche to the boundary, R is the plate radius, and k = 1000. *ϵ* regulates the shape of the impact function. We deducted the influence from the colony after each discrete time step. For the purpose of encryption, we maximized the influence of the geometry by modulating *ϵ*, such that the negative plate impact reached as far as the center of the patterns. We used *ϵ* = 1 for generating the encryption datasets, and 2000 for any other dataset.

When using the shape of the growing medium as the secret key, we simulated the colony patterns on circular, diamond, square and equilateral triangular shaped domains. The area of each geometry was kept the same in order to compare the effect of the geometry. We removed the information of growth domain shape by cropping out a smaller, circular area at the center of each pattern, only the processed pattern images were used for CNN training.

### Ensemble Learning and Uncertainty Estimation

The training of ensemble model was carried out in two steps. First, we trained several base CNN models using the same protocol described in “CNN training”. Their probabilistic predictions on the training set were then linearly combined to constitute a new dataset. Next, we used the new dataset to train an ensemble model from scratch. We tested several ensemble model architectures, including logistic regression and feedforward neural networks with different number of hidden layers and nodes. In the ensemble model, we used ReLU activation function for the input and hidden layers and passed the model output into softmax function to turn it into categorical probabilities. For its training, we used Glorot uniform initializer, categorical cross entropy loss and Adam optimization algorithm with learning rate = 0.0001. Keras early stopping was used to stop the training once the loss metric stopped improving. The patience was 5 and the minimum change was 0. 0001. We evaluated the model performance on a balanced dataset of 100 datapoints per class through metrics such as precision, recall, ROC, AUC ROC using scikit-learn (0.22.2).

We evaluated the prediction uncertainty based on the output of base models. We used common metrics, such as log likelihood, mean square error (MSE), top-1 and top-5 errors, for estimating the uncertainty. Specifically, the log likelihood is 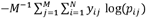 and the MSE is 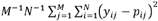. For the i^th^ data point, *y*_*ij*_ is the true label for class j (1 if the data point belongs to class j, otherwise 0), *p*_*ij*_ is the predicted probabilities for class j. M indicates the total number of data points, and N indicates the dictionary size. Top-1 and top 5 the fraction of data points whose correct label is not among their top 1 or 5 probable predictions, respectively.

## Data and Code Availability

The mathematical simulation and machine learning codes used in this study are available on GitHub: https://github.com/youlab/Information_encoding. The platform for encoding text in the format of video is available at https://www.patternencoder.com/.

## Acknowledgments

We thank Thomas Witelski, Sayan Mukherjee and Teng Wang for the helpful discussion and comments on the manuscript. We thank Duke Compute Cluster for assistance with high-throughput computation. This study was partially supported by the National Science Foundation (MCB-1937259), the Office of Naval Research (N00014-20-1-2121), the David and Lucile Packard Foundation, and the Google Cloud Research Credits program.

