## Appendix A for "Distributed information encoding and decoding using self-organized spatial patterns"

**Encoding characters in Emorfi.**

To construct Emorfi, each of the printable ASCII characters (including English letters in upper and lower cases, digits, punctuations, and whitespaces) was converted into a binary representation, which was then converted into a unique initial configuration. For each configuration, we carried out mathematical simulation and obtained 1000 different patterns. These 100 sets of patterns, as well as subsequent ones to be generated, make up Emorfi. In the table below, three examples are shown for each character.

| Encoding character | Binary representation | Initial configuration | Example patterns | | |
| --- | --- | --- | --- | --- | --- |
| 0 | 0000001 | 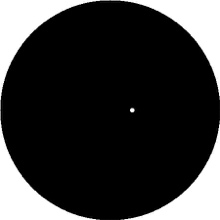 | 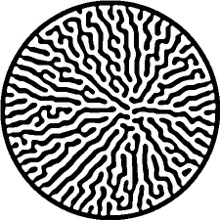 | 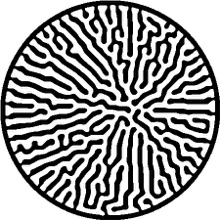 | 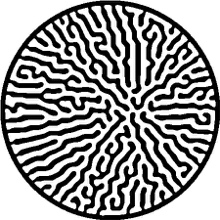 |
| 1 | 0000010 | 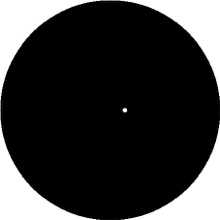 | 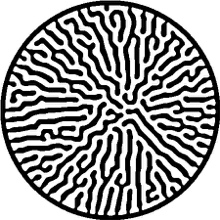 | 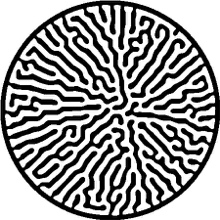 | 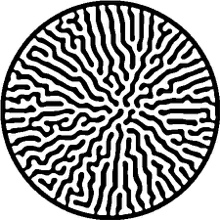 |
| 2 | 0000011 | 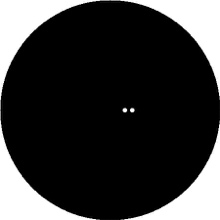 | 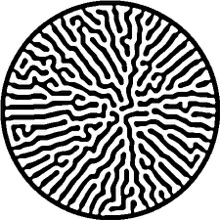 | 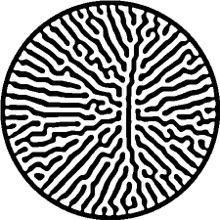 | 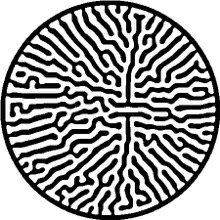 |
| 3 | 0000100 | 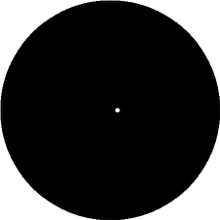 | 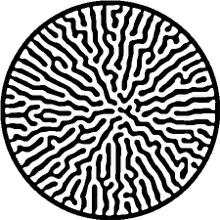 | 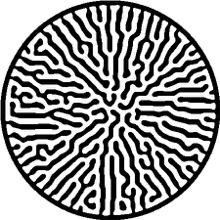 | 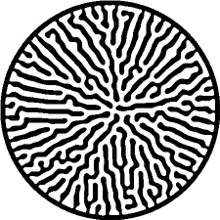 |
| 4 | 0000101 | 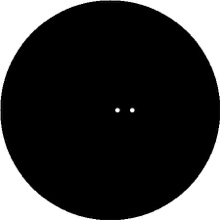 | 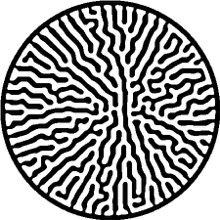 | 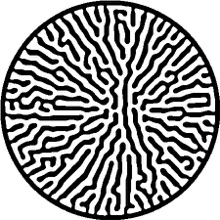 | 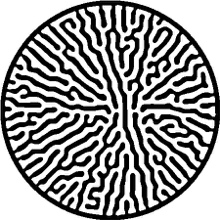 |
| 5 | 0000110 | 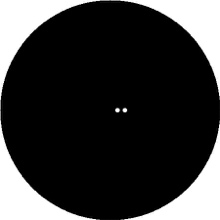 | 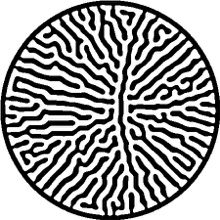 | 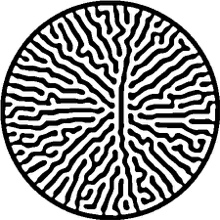 | 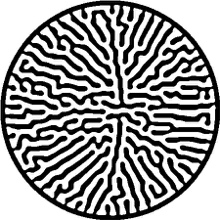 |
| 6 | 0000111 | 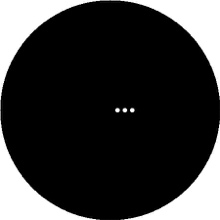 | 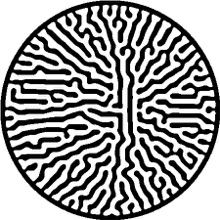 | 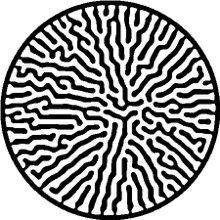 | 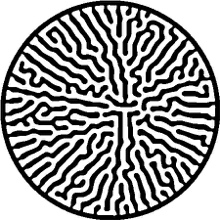 |
| 7 | 0001000 | 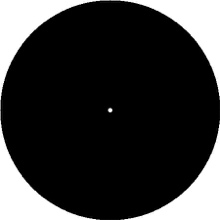 | 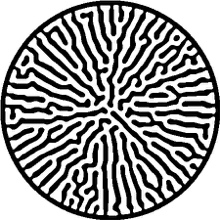 | 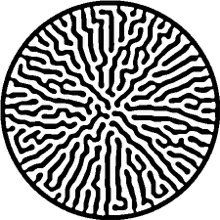 | 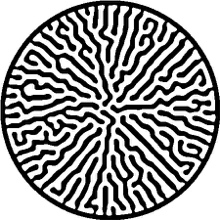 |
| 8 | 0001001 | 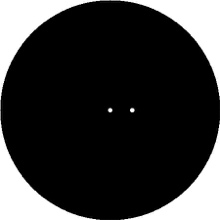 | 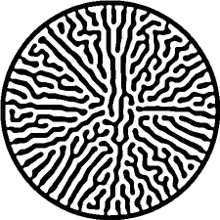 | 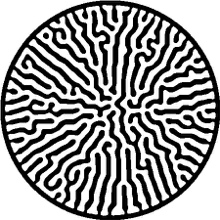 | 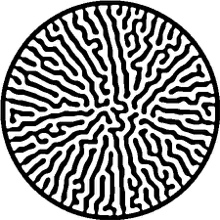 |
| 9 | 0001010 | 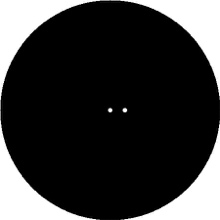 | 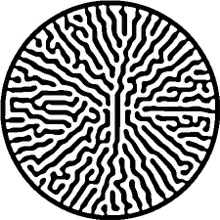 | 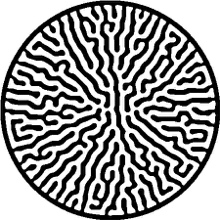 | 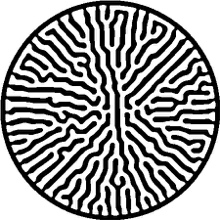 |
| a | 0001011 | 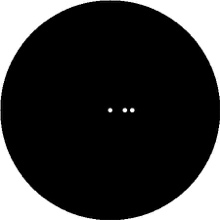 | 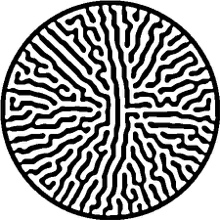 | 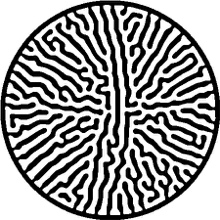 | 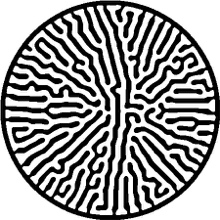 |
| b | 0001100 | 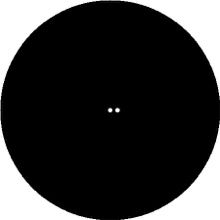 | 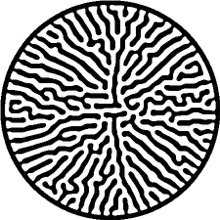 | 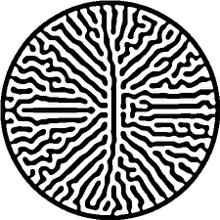 | 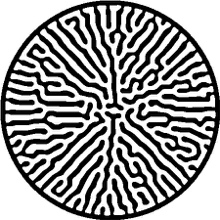 |
| c | 0001101 | 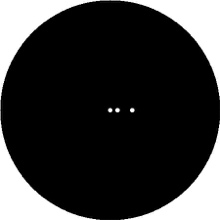 | 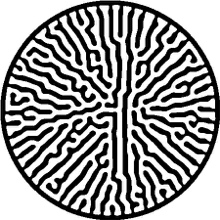 | 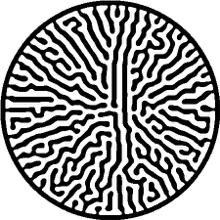 | 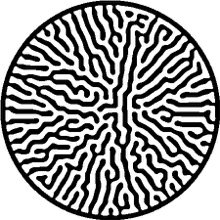 |
| d | 0001110 | 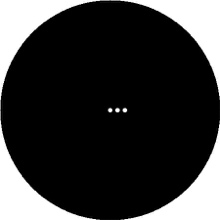 | 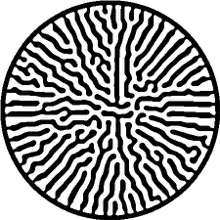 | 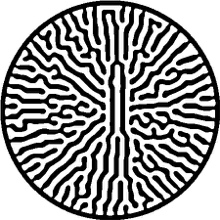 | 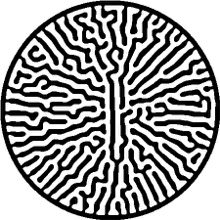 |
| e | 0001111 | 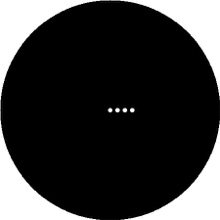 | 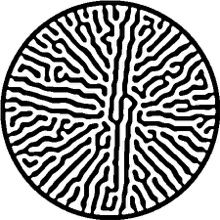 | 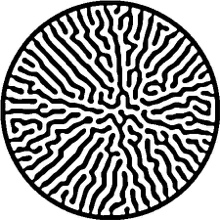 | 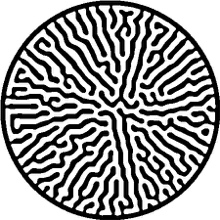 |
| f | 0010000 | 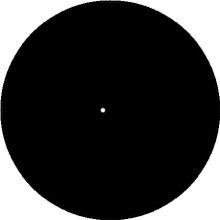 | 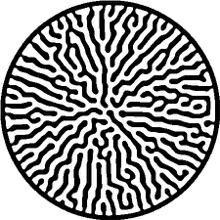 | 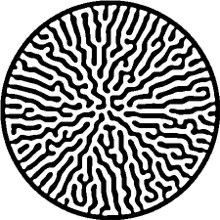 | 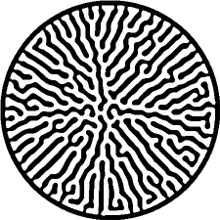 |
| g | 0010001 | 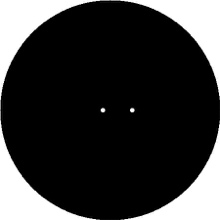 | 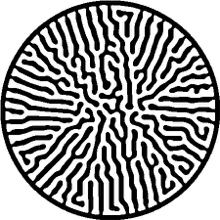 | 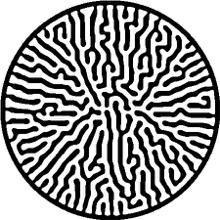 | 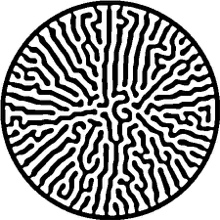 |
| h | 0010010 | 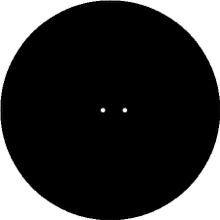 | 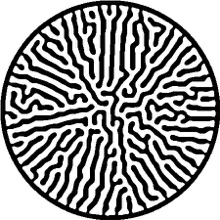 | 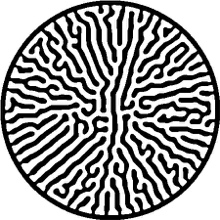 | 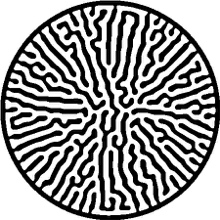 |
| i | 0010011 | 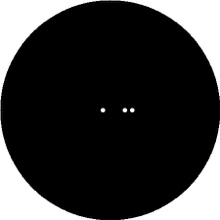 | 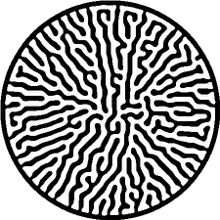 | 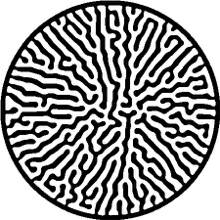 | 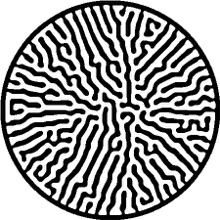 |
| j | 0010100 | 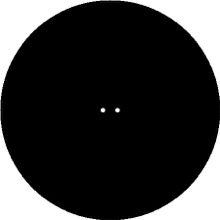 | 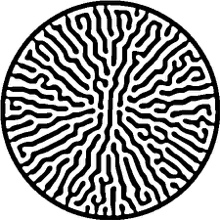 | 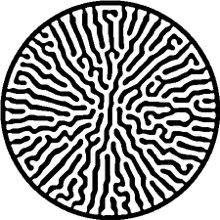 | 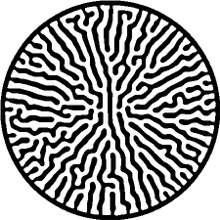 |
| k | 0010101 | 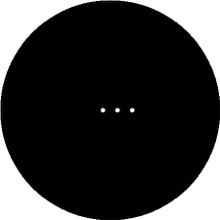 | 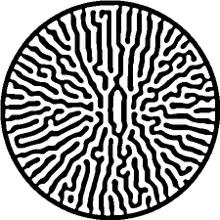 | 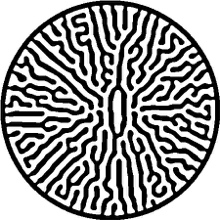 | 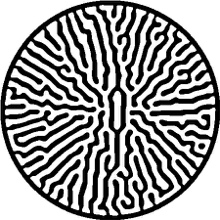 |
| l | 0010110 | 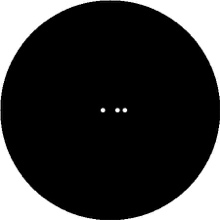 | 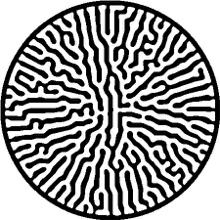 | 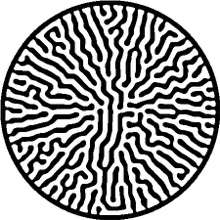 | 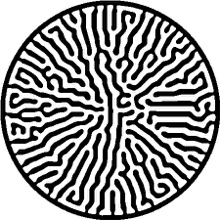 |
| m | 0010111 | 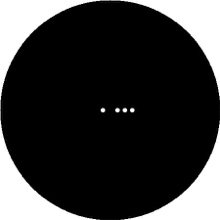 | 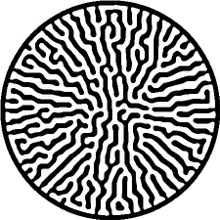 | 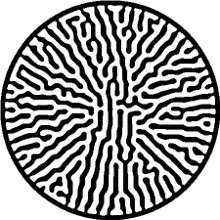 | 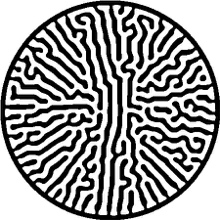 |
| n | 0011000 | 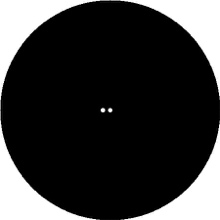 | 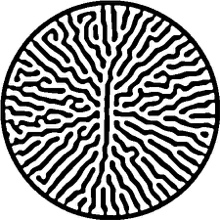 | 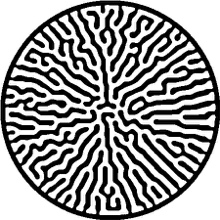 | 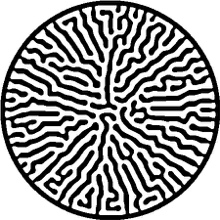 |
| o | 0011001 | 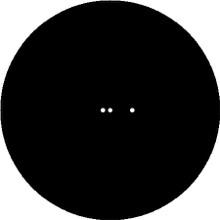 | 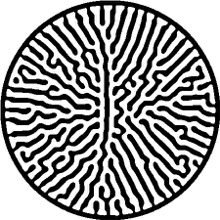 | 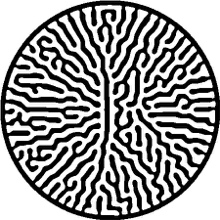 | 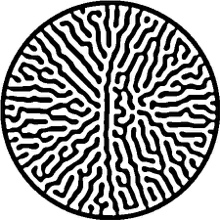 |
| p | 0011010 |  |  |  |  |
| q | 0011011 |  |  |  |  |
| r | 0011100 |  |  |  |  |
| s | 0011101 |  |  |  |  |
| t | 0011110 |  |  |  |  |
| u | 0011111 |  |  |  |  |
| v | 0100000 |  |  |  |  |
| w | 0100001 |  |  |  |  |
| x | 0100010 |  |  |  |  |
| y | 0100011 |  |  |  |  |
| z | 0100100 |  |  |  |  |
| A | 0100101 |  |  |  |  |
| B | 0100110 |  |  |  |  |
| C | 0100111 |  |  |  |  |
| D | 0101000 |  |  |  |  |
| E | 0101001 |  |  |  |  |
| F | 0101010 |  |  |  |  |
| G | 0101011 |  |  |  |  |
| H | 0101100 |  |  |  |  |
| I | 0101101 |  |  |  |  |
| J | 0101110 |  |  |  |  |
| K | 0101111 |  |  |  |  |
| L | 0110000 |  |  |  |  |
| M | 0110001 |  |  |  |  |
| N | 0110010 |  |  |  |  |
| O | 0110011 |  |  |  |  |
| P | 0110100 |  |  |  |  |
| Q | 0110101 |  |  |  |  |
| R | 0110110 |  |  |  |  |
| S | 0110111 |  |  |  |  |
| T | 0111000 |  |  |  |  |
| U | 0111001 |  |  |  |  |
| V | 0111010 |  |  |  |  |
| W | 0111011 |  |  |  |  |
| X | 0111100 |  |  |  |  |
| Y | 0111101 |  |  |  |  |
| Z | 0111110 |  |  |  |  |
| ！ | 0111111 |  |  |  |  |
| “ | 1000000 |  |  |  |  |
| # | 1000001 |  |  |  |  |
| $ | 1000010 |  |  |  |  |
| % | 1000011 |  |  |  |  |
| & | 1000100 |  |  |  |  |
| ‘ | 1000101 |  |  |  |  |
| ( | 1000110 |  |  |  |  |
| ) | 1000111 |  |  |  |  |
| * | 1001000 |  |  |  |  |
| + | 1001001 |  |  |  |  |
| ， | 1001010 |  |  |  |  |
| - | 1001011 |  |  |  |  |
| . | 1001100 |  |  |  |  |
| / | 1001101 |  |  |  |  |
| : | 1001110 |  |  |  |  |
| ; | 1001111 |  |  |  |  |
| < | 1010000 |  |  |  |  |
| = | 1010001 |  |  |  |  |
| > | 1010010 |  |  |  |  |
| ? | 1010011 |  |  |  |  |
| @ | 1010100 |  |  |  |  |
| [ | 1010101 |  |  |  |  |
| \\ | 1010110 |  |  |  |  |
| ] | 1010111 |  |  |  |  |
| ^ | 1011000 |  |  |  |  |
| _ | 1011001 |  |  |  |  |
| ` | 1011010 |  |  |  |  |
| { | 1011011 |  |  |  |  |
| ｜ | 1011100 |  |  |  |  |
| } | 1011101 |  |  |  |  |
| ~ | 1011110 |  |  |  |  |
| Character space | 1011111 |  |  |  |  |
| \t (tab) | 1100000 |  |  |  |  |
| \n (linefeed) | 1100001 |  |  |  |  |
| \r (return) | 1100010 |  |  |  |  |
| \x0b (vertical tab) | 1100011 |  |  |  |  |
| \x0c (formfeed) | 1100100 |  |  |  |  |
