## Appendix B for "Distributed information encoding and decoding using self-organized spatial patterns"

**Encoding “I Have a Dream” in Emorfi and decoding it back in English.**

Each character was encoded in a randomly selected pattern from its corresponding class (see Appendix A). Next, the patterns were assembled in order into a video. To accommodate majority voting, the above process was repeated four times and the new video was concatenated to the end of the existing one. When decoding, we used a trained CNN combined with majority voting, which led to a decoding accuracy of 99.8%.

**I Have a Dream**

**By Martin Luther King Jr.**

**August 28, 1963**

**Original:**

Five score years ago, a great American, in whose symbolic shadow we stand today, signed the Emancipation Proclamation. This momentous decree came as a great beacon light of hope to millions of Negro slaves who had been seared in the flames of withering injustice. It came as a joyous daybreak to end the long night of their captivity. But 100 years later, the Negro still is not free. One hundred years later, the life of the Negro is still sadly crippled by the manacles of segregation and the chains of discrimination. One hundred years later, the Negro lives on a lonely island of poverty in the midst of a vast ocean of material prosperity. One hundred years later the Negro is still languished in the corners of American society and finds himself in exile in his own land. And so we've come here today to dramatize a shameful condition. In a sense we've come to our nation's capital to cash a check. When the architects of our republic wrote the magnificent words of the Constitution and the Declaration of Independence, they were signing a promissory note to which every American was to fall heir. This note was a promise that all men - yes, black men as well as white men - would be guaranteed the unalienable rights of life, liberty and the pursuit of happiness. It is obvious today that America has defaulted on this promissory note insofar as her citizens of color are concerned. Instead of honoring this sacred obligation, America has given the Negro people a bad check, a check which has come back marked insufficient funds. But we refuse to believe that the bank of justice is bankrupt. We refuse to believe that there are insufficient funds in the great vaults of opportunity of this nation. And so we've come to cash this check, a check that will give us upon demand the riches of freedom and the security of justice. We have also come to his hallowed spot to remind America of the fierce urgency of now. This is no time to engage in the luxury of cooling off or to take the tranquilizing drug of gradualism. Now is the time to make real the promises of democracy. Now is the time to rise from the dark and desolate valley of segregation to the sunlit path of racial justice. Now is the time to lift our nation from the quick sands of racial injustice to the solid rock of brotherhood. Now is the time to make justice a reality for all of God's children. It would be fatal for the nation to overlook the urgency of the moment. This sweltering summer of the Negro's legitimate discontent will not pass until there is an invigorating autumn of freedom and equality. 1963 is not an end, but a beginning. Those who hope that the Negro needed to blow off steam and will now be content will have a rude awakening if the nation returns to business as usual. There will be neither rest nor tranquility in America until the Negro is granted his citizenship rights. The whirlwinds of revolt will continue to shake the foundations of our nation until the bright day of justice emerges. But there is something that I must say to my people who stand on the warm threshold which leads into the palace of justice. In the process of gaining our rightful place, we must not be guilty of wrongful deeds. Let us not seek to satisfy our thirst for freedom by drinking from the cup of bitterness and hatred. We must forever conduct our struggle on the high plane of dignity and discipline. We must not allow our creative protest to degenerate into physical violence. Again and again, we must rise to the majestic heights of meeting physical force with soul force. The marvelous new militancy which has engulfed the Negro community must not lead us to a distrust of all white people, for many of our white brothers, as evidenced by their presence here today, have come to realize that their destiny is tied up with our destiny. And they have come to realize that their freedom is inextricably bound to our freedom. We cannot walk alone. And as we walk, we must make the pledge that we shall always march ahead. We cannot turn back. There are those who are asking the devotees of civil rights, when will you be satisfied? We can never be satisfied as long as the Negro is the victim of the unspeakable horrors of police brutality. We can never be satisfied as long as our bodies, heavy with the fatigue of travel, cannot gain lodging in the motels of the highways and the hotels of the cities. We cannot be satisfied as long as the Negro's basic mobility is from a smaller ghetto to a larger one. We can never be satisfied as long as our children are stripped of their selfhood and robbed of their dignity by signs stating: for whites only. We cannot be satisfied as long as a Negro in Mississippi cannot vote and a Negro in New York believes he has nothing for which to vote. No, no, we are not satisfied, and we will not be satisfied until justice rolls down like waters, and righteousness like a mighty stream. I am not unmindful that some of you have come here out of great trials and tribulations. Some of you have come fresh from narrow jail cells. Some of you have come from areas where your quest for freedom left you battered by the storms of persecution and staggered by the winds of police brutality. You have been the veterans of creative suffering. Continue to work with the faith that unearned suffering is redemptive. Go back to Mississippi, go back to Alabama, go back to South Carolina, go back to Georgia, go back to Louisiana, go back to the slums and ghettos of our Northern cities, knowing that somehow this situation can and will be changed. Let us not wallow in the valley of despair, I say to you today, my friends. So even though we face the difficulties of today and tomorrow, I still have a dream. It is a dream deeply rooted in the American dream. I have a dream that one day this nation will rise up and live out the true meaning of its creed: We hold these truths to be self-evident, that all men are created equal. I have a dream that one day on the red hills of Georgia, the sons of former slaves and the sons of former slave owners will be able to sit down together at the table of brotherhood. I have a dream that one day even the state of Mississippi, a state sweltering with the heat of injustice, sweltering with the heat of oppression will be transformed into an oasis of freedom and justice. I have a dream that my four little children will one day live in a nation where they will not be judged by the color of their skin but by the content of their character. I have a dream today. I have a dream that one day down in Alabama with its vicious racists, with its governor having his lips dripping with the words of interposition and nullification, one day right down in Alabama little black boys and black girls will be able to join hands with little white boys and white girls as sisters and brothers. I have a dream today. I have a dream that one day every valley shall be exalted, every hill and mountain shall be made low, the rough places will be made plain, and the crooked places will be made straight, and the glory of the Lord shall be revealed, and all flesh shall see it together. This is our hope. This is the faith that I go back to the South with. With this faith, we will be able to hew out of the mountain of despair a stone of hope. With this faith we will be able to transform the jangling discords of our nation into a beautiful symphony of brotherhood. With this faith we will be able to work together, to pray together, to struggle together, to go to jail together, to stand up for freedom together, knowing that we will be free one day. This will be the day when all of God's children will be able to sing with new meaning: My country, 'tis of thee, sweet land of liberty, of thee I sing. Land where my fathers died, land of the pilgrims' pride, from every mountainside, let freedom ring. And if America is to be a great nation, this must become true. And so let freedom ring from the prodigious hilltops of New Hampshire. Let freedom ring from the mighty mountains of New York. Let freedom ring from the heightening Alleghenies of Pennsylvania. Let freedom ring from the snowcapped Rockies of Colorado. Let freedom ring from the curvaceous slopes of California. But not only that, let freedom ring from Stone Mountain of Georgia. Let freedom ring from Lookout Mountain of Tennessee. Let freedom ring from every hill and molehill of Mississippi. From every mountainside, let freedom ring. And when this happens, and when we allow freedom ring, when we let it ring from every village and every hamlet, from every state and every city, we will be able to speed up that day when all of God's children, black men and white men, Jews and Gentiles, Protestants and Catholics, will be able to join hands and sing in the words of the old Negro spiritual: Free at last. Free at last. Thank God almighty, we are free at last.

**Decoded:** (the decoding accuracy is 99.8 %, the mispredicted characters are labeled in red)

Five score years ago, a great American, in whose symbolic shadow we stand today, signed the Emancipation Proclamation. This momentous decree came as a great beacon light of hope to millions of Negro slaves who had been seared in the flames of withering injustice. It came as a joyous daybreak to end the long night of their captivity. But 100 years later, the Negro still is not free. One hundred years later, the life of the Negro is still sadly crippled by the manacles of segregation and the chains of discrimination. One hundred years later, the Negro lives on a lonely island of poverty in the midst of a vast ocean of material prosperity. One hundred years later the Negro is still languished in the corners of American society and finds himself in exile in his own land. And so we've come here today to dramatize a shameful condition. In a sense we've come to our nation's capital to cash a check. When the architects of our republic wrote the magnificent words of the Constitution and the Declaration of Independence, they were signing a promissory note to which every American was to fall heir. This note was a promise that all men - yes, black men as well as white men - would be guaranteed the unalienable rights of life, liberty and the pursuit of happiness. It is obvious today that America has default**c**d on this promissory note insofar as her citizens of color are concerned. Instead of honoring this sacred obligation, America has given the Negro people a bad check, a check which has come back marked insufficient funds. But we refuse to believe that the bank of justice is bankrupt. We refuse to believe that there are insufficient funds in the great vaults of opportunity of this nation. And so we've come to cash this check, a check that will give us upon demand the riches of freedom and the security of justice. We have also come to his hallowed spot to remind America of the fierce urgency of now. This is no time to engage in the luxury of cooling off or to take the tranquilizing drug of gradualism. Now is the time to make real the promises of democracy. Now is the time to rise from the dark and desolate valley of segregation to the sunlit path of racial justice. Now is the time to lift our nation from the quick sands of racial injustice to the solid rock of brotherhood. Now is the time to make justice a reality for all of God's children. It would be fatal for the nation to overlook the urgency of the moment. This sweltering summer of the Negro's legitimate discontent will not pass until there is an invigorating autumn of freedom and equality. 1963 is not an end, but a beginning. Those who hope that the Negro needed to blow off steam and will now be content will have a rude awakening if the nation returns to business as usual. There will be neither rest nor tranquility in America until the Negro is granted his citizenship rights. The whirlwinds of revolt will continue to shake the foundations of our nation until the bright day of justice emerges. But there is something that I must say to my people who stand on the warm threshold which leads into the palace of justice. In the process of gaining our rightful place, we must not be guilty of wrongful de**8**ds. Let us not seek to satisfy our thirst for freedom by drinking from the cup of bitterness and hatred. We must forever conduct our struggle on the high plane of dignity and discipline. We must not allow our creative protest to degenerate into physical violence. Again and again, we must rise to the majestic heights of meeting physical force with soul force. The marvelous new militancy which has engulfed the Negro community must not lead us to a distrust of all white people, for many of our white brothers, as evidenced by their presence here today, have come to realize that their destiny is tied up with our destiny. And they have come to realize **u**hat their freedom is inextricably bound to our freedom. We cannot walk alone. And as we walk, we must make the pledge that we shall always march ahead. We cannot turn back. There are those who are asking the devotees of civil rights, when will you be satisfied? We can never be satisfied as long as the Negro is the victim of the unspeakable horrors of police brutality. We can never be satisfied as**{**long as our bodies, heavy with the fatigue of travel, cannot gain lodging in the motels of the highways and the hotels of th**d** cities. We cannot be satisfied as long as the Negro's basic mobility is from a smaller ghetto to a larger one. We can nev**q**r be satisfied as long as our children are stripped of their selfhood and robbed of their dignity by signs stating: for whites only. We cannot be satisfied as long as a Negro in Mississippi cannot vote and a Negro in New York believes he has nothing for which to vote. No, no, we are not satisfied, and we will not be satisfied until justice rolls down like waters, and righteousness like a mighty stream. I am not unmindful that some of you have come here out of great trials and tribulations. Some of you have come fresh from narrow jail cells. Some of you have come from areas where your quest for freedom left you battered by the storms of persecution and staggered by the winds of police brutality. You have been the veterans of creative suffering. Continue to work with the faith that unearned suffering is redemptive. Go back to Mississippi, go back to Alabama, go back to South Carolina, go back to Georgia, go back to Louisiana, go back to the slums and ghettos of our Northern cities, knowing that somehow this situation can and will be changed. Let us not wallow in**{**the valley of despair, I say to you today, my friends. So even though we face the difficulties of today and tomorrow, I still have a dream. It is a dream deeply rooted in the American dream. I have a dream that one day this nation will rise up and live out the true meaning of its creed:**{**We hold these truths to be self-evident, that all men are created equal. I have a dream that one day on the red hills of Georgia, the sons of former slaves and the sons of former slave owners will be able to sit**{**down together at the table of brotherhood. I have a dream that one day even the state of Mississippi, a state sweltering with the heat of injustice, sweltering with the heat of oppression will be transformed into an oasis of freedom and justice. I have a dream that my four little children will one day live in a nation where they will not be judged by the color of their skin but by the content of their character. I have a dream today. I have a dream that one day down in Alabama with its vicious racists, with its governor having his lips dripping with the words of interposition and**~**nullification, one day right down in Alabama little black boys and black girls will be able to join hands with little white boys and white girls as sisters and brothers. I have a dream today. I have a dr**u**am that one day every valley shall be exalted, every hill and mountain shall b**8** made**{**low, the rough places will be made plain, and the crooked places will be made straight, and the glory of the Lord shall be revealed, and all flesh shall see it together. This is our hope. This is the faith that I go**_**back to the South with. With this faith, we will be able to hew out of the mountain of despair a stone of hope. With this faith we will be able to transform the jangling discords of our nation into a beautiful symphony of brotherhood. With this faith we will**{**be able to work together, to pray together, to struggle together, to go to jail together, to stand up for freedom together, knowing that we will be free one day. This will be the day when all of God's children will be able to sing with new meaning: My country, 'tis of thee, sweet land of liberty, of thee I sing. Land where my fathers died, land of the pilgrims' pride, from every mountainside, let freedom ring. And if America is to be a great nation, this must become true. And so let freedom ring from the prodigious hilltops of New Hampshire. Let freedom ring from the mighty mountains of New York. Let freedom ring from the heightening Alleghenies of Pennsylvania. Let freedom ring from the snowcapped Rockies of Colorado. Let freedom ring from the curvaceous slopes of California. But not only that, let freedom ring from Stone Mountain of Georgia. Let freedom ring from Lookout Mountain of Tennessee. Let freedom ring from every hill**{**and molehill of Mississippi. From every mountainside, let freedom ring. And when this happens, and when we allow freedom ring, when we let it ring from every village and every hamlet, from every state and every city, we will be able to speed up that day when all of God's children, black men and white men, Jews and Gentiles, Protestants and Catholics, will be able to join hands and sing in the words of the old N**c**gro spiritual: Fr**8**e at last. Free at last. Thank God almighty, we are free at last.
