## Appendix C for "Distributed information encoding and decoding using self-organized spatial patterns"

**Encoding “Auguries of Innocence” in Emorfi and decoding it back in English.**

Each character was encoded in a randomly selected pattern from its corresponding class (see Appendix A). Next, the patterns were assembled in order into a video. To accommodate majority voting, the above process was repeated four times and the new video was concatenated to the end of the existing one. When decoding, we used a trained CNN combined with majority voting, which led to a decoding accuracy of 99.6%.

**Auguries of Innocence**

**By William Blake**

**1863**

**Original** (Poetry foundation):

To see a World in a Grain of Sand

And a Heaven in a Wild Flower

Hold Infinity in the palm of your hand

And Eternity in an hour

A Robin Red breast in a Cage

Puts all Heaven in a Rage

A Dove house filld with Doves & Pigeons

Shudders Hell thr' all its regions

A dog starvd at his Masters Gate

Predicts the ruin of the State

A Horse misusd upon the Road

Calls to Heaven for Human blood

Each outcry of the hunted Hare

A fibre from the Brain does tear

A Skylark wounded in the wing

A Cherubim does cease to sing

The Game Cock clipd & armd for fight

Does the Rising Sun affright

Every Wolfs & Lions howl

Raises from Hell a Human Soul

The wild deer, wandring here & there

Keeps the Human Soul from Care

The Lamb misusd breeds Public Strife

And yet forgives the Butchers knife

The Bat that flits at close of Eve

Has left the Brain that wont Believe

The Owl that calls upon the Night

Speaks the Unbelievers fright

He who shall hurt the little Wren

Shall never be belovd by Men

He who the Ox to wrath has movd

Shall never be by Woman lovd

The wanton Boy that kills the Fly

Shall feel the Spiders enmity

He who torments the Chafers Sprite

Weaves a Bower in endless Night

The Catterpiller on the Leaf

Repeats to thee thy Mothers grief

Kill not the Moth nor Butterfly

For the Last Judgment draweth nigh

He who shall train the Horse to War

Shall never pass the Polar Bar

The Beggars Dog & Widows Cat

Feed them & thou wilt grow fat

The Gnat that sings his Summers Song

Poison gets from Slanders tongue

The poison of the Snake & Newt

Is the sweat of Envys Foot

The poison of the Honey Bee

Is the Artists Jealousy

The Princes Robes & Beggars Rags

Are Toadstools on the Misers Bags

A Truth thats told with bad intent

Beats all the Lies you can invent

It is right it should be so

Man was made for Joy & Woe

And when this we rightly know

Thro the World we safely go

Joy & Woe are woven fine

A Clothing for the soul divine

Under every grief & pine

Runs a joy with silken twine

The Babe is more than swadling Bands

Throughout all these Human Lands

Tools were made & Born were hands

Every Farmer Understands

Every Tear from Every Eye

Becomes a Babe in Eternity

This is caught by Females bright

And returnd to its own delight

The Bleat the Bark Bellow & Roar

Are Waves that Beat on Heavens Shore

The Babe that weeps the Rod beneath

Writes Revenge in realms of Death

The Beggars Rags fluttering in Air

Does to Rags the Heavens tear

The Soldier armd with Sword & Gun

Palsied strikes the Summers Sun

The poor Mans Farthing is worth more

Than all the Gold on Africs Shore

One Mite wrung from the Labrers hands

Shall buy & sell the Misers Lands

Or if protected from on high

Does that whole Nation sell & buy

He who mocks the Infants Faith

Shall be mockd in Age & Death

He who shall teach the Child to Doubt

The rotting Grave shall neer get out

He who respects the Infants faith

Triumphs over Hell & Death

The Childs Toys & the Old Mans Reasons

Are the Fruits of the Two seasons

The Questioner who sits so sly

Shall never know how to Reply

He who replies to words of Doubt

Doth put the Light of Knowledge out

The Strongest Poison ever known

Came from Caesars Laurel Crown

Nought can Deform the Human Race

Like to the Armours iron brace

When Gold & Gems adorn the Plow

To peaceful Arts shall Envy Bow

A Riddle or the Crickets Cry

Is to Doubt a fit Reply

The Emmets Inch & Eagles Mile

Make Lame Philosophy to smile

He who Doubts from what he sees

Will neer Believe do what you Please

If the Sun & Moon should Doubt

Theyd immediately Go out

To be in a Passion you Good may Do

But no Good if a Passion is in you

The Whore & Gambler by the State

Licencd build that Nations Fate

The Harlots cry from Street to Street

Shall weave Old Englands winding Sheet

The Winners Shout the Losers Curse

Dance before dead Englands Hearse

Every Night & every Morn

Some to Misery are Born

Every Morn and every Night

Some are Born to sweet delight

Some are Born to sweet delight

Some are Born to Endless Night

We are led to Believe a Lie

When we see not Thro the Eye

Which was Born in a Night to perish in a Night

When the Soul Slept in Beams of Light

God Appears & God is Light

To those poor Souls who dwell in Night

But does a Human Form Display

To those who Dwell in Realms of day

**Decoded**: (the decoding accuracy is 99.6%, the mispredicted characters are labeled in red)

To see a World in a Grain of Sand

And a Heaven in a Wild Flower

Hold Infinity in the palm of your hand

And Eternity in an hour

A Robin Red breast in a Cage

Puts all Heaven*****in a Rage

A Dove house filld with Doves & Pigeons

Shudders Hell thr' all its regions

A dog starvd at his Masters Gate

Predicts the ruin of the State

A Horse misusd upon the Road

Calls to Heaven for Human blood

Each outcry of the hunted Hare

A fibre from the Brain does tear

A Skylark wounded in the wing

A Cherubim does cease to sing

The Game Cock clipd & armd for fight

Does the Rising Sun affright

Every Wolfs & Lions howl

Raises from Hell a Human Soul

The wild deer, wandring here & there

Keeps the Human Soul from Care

The Lamb misusd breeds Public Strife

And yet forgives the Butchers knife

The Bat that flit**u** at close of Eve

Has left the Brain that wont Believe

The Owl t**l**at calls upon the Night

Speaks the Unbelievers fright

He who shall hurt the little Wren

Shall never be belovd by Men

He who the Ox to**{**wrath has movd

Shall never be by Woman lovd

The wanton Boy that kills the Fly

Shall feel the Spiders enmity

He who torments the Chafers Sprite

Weaves a Bower in endless Night

The Catterpiller on the Leaf

Repeats to thee thy Mothers grief

Kill not the Moth nor Butterfly

For the Last Judgment draweth nigh

He who shall train the Horse to War

Shall never pass the Polar Bar

The Beggars Dog & Widows Cat

Feed them & thou wilt grow fat

The Gnat that sings his Summers Song

Poison gets from Slanders tongue

The poison of the Snake & Newt

Is the sweat of Envys Foot

The poison of the Honey Bee

Is the Artists Jealousy

The Princes Robes & Beggars Rags

Are Toadstools on the Misers Bags

A Truth thats told with bad intent

Beats all the Lies you can invent

It is right it should be so

Man was made for Joy & Woe

And when this we rightly know

Thro the World we safely go

Joy & Woe are woven fine

A Clothing for the**?**soul divine

Under every grief & pine

Runs a joy with silken twine

The Babe is more than swadling Bands

Throughout all these Human Lands

Tools were made &**]**Born were hands

Every Farmer Understands

Every Tear from Every Eye

Becomes a Babe in Eternity

This is caught by Female**u** bright

And returnd to its own delight

The Bleat the Bark Bellow & Roar

Are Waves that Beat on Heavens Shore

The Babe**~**that weeps the Rod beneath

Writes Revenge in realms**{**of Death

The Beggars Rags fluttering in Air

Does to Rags the Heavens tear

The Soldier armd with Sword & Gun

Palsied strikes the Summers Sun

The poor Mans Farthing is worth more

Than all the Gold on Africs Shore

One Mite wrung from the Labrers hands

Shall buy & sell the Misers Lands

Or if p**t**otected from on high

Does that whole Nation sell & buy

He who mocks the Infants Faith

Shall be mockd in Ag**c** & Death

He who shall teach the Child to Doubt

The rotting Grave shall neer get out

He who respects the Infants faith

Triumphs over Hell & Death

The Childs Toys & the Old Mans Reasons

Are the Fruits of the Two seasons

The Questioner who sits**?**so sly

Shall never know how to Reply

He who replies to words of Doubt

Doth put the Light of Knowledge out

The Strongest Poison ever known

Came from Caesars Laurel**~**Crown

Nought can Deform the Human Race

Like to the Armours iron brace

When Gold & Gems adorn the Plow

To peaceful Arts shall Envy Bow

A Riddle or the Crickets Cry

Is to Doubt a fit Reply

The Emmets Inch & Eagles Mile

Make Lame Philosophy to smile

He**{**who Doubts from what he sees

Will neer Believe do what you Please

If**]**the Sun & Moon should Doubt

Theyd immediately Go out

To be in a Passion you Good may Do

But no Good if a Passion is in you

The Whore & Gambler by the State

Licencd build that Nations Fate

The Harlots cry from Street to Street

Shall weave Old Englands winding Sheet

The Winners Shout the Losers Curse

Dance before dead Englands Hearse

Every Night & every Morn

Some to Misery are Born

Every Morn and every Night

Some are Born to sweet delight

Some are Born to sweet delight

Some are Born to Endless Night

We are led to Believe a Lie

When we see not Thro the Eye

Which was Born in a Night to perish in a Night

When the Soul Sl**c**pt in Beams of Light

God Appears & God is Light

To those poor Souls who dwell in Night

But does a Human Form Display

To those who Dwell in Realms of day
