## Appendix D for "Distributed information encoding and decoding using self-organized spatial patterns"

**Encoding the GFP protein sequence in patterns and decoding it back in amino acid.**

A similar approach for generating Emorfi was used to encode and decode protein sequences. Each of the 20 common amino acids was converted into a binary representation and then converted into a unique initial configuration. 1000 images were generated through mathematical simulation for each amino acid.

To encode GFP sequence, each amino acid was encoded in a randomly selected pattern from its corresponding class. Next, the patterns were assembled in order into a video. To accommodate majority voting, the above process was repeated four times and the new video was concatenated to the end of the existing one. When decoding, we used a trained CNN combined with majority voting, which led to a decoding accuracy of 100%.

**GFP protein sequence**

**Original** (Uniprot):

MSKGEELFTGVVPILVELDGDVNGHKFSVSGEGEGDATYGKLTLKFICTTGKLPVPWPTLVTTFSYGVQCFSRYPDHMKQHDFFKSAMPEGYVQERTIFFKDDGNYKTRAEVKFEGDTLVNRIELKGIDFKEDGNILGHKLEYNYNSHNVYIMADKQKNGIKVNFKIRHNIEDGSVQLADHYQQNTPIGDGPVLLPDNHYLSTQSALSKDPNEKRDHMVLLEFVTAAGITHGMDELYK

**Decoded:** (the decoding accuracy is 100%)

MSKGEELFTGVVPILVELDGDVNGHKFSVSGEGEGDATYGKLTLKFICTTGKLPVPWPTLVTTFSYGVQCFSRYPDHMKQHDFFKSAMPEGYVQERTIFFKDDGNYKTRAEVKFEGDTLVNRIELKGIDFKEDGNILGHKLEYNYNSHNVYIMADKQKNGIKVNFKIRHNIEDGSVQLADHYQQNTPIGDGPVLLPDNHYLSTQSALSKDPNEKRDHMVLLEFVTAAGITHGMDELYK
