## Supplementary Information for "Distributed information encoding and decoding using self-organized spatial patterns"

Supplementary Information Text

**Temporal information encoding and decoding**

In dynamical systems, information embedded in the initial conditions may dissipate over time(1). Thus, it is necessary to determine the temporal encoding-decoding performance of our patterning system. We simulated branching patterns using the same protocol as described in the Methods. Instead of waiting until the colony stops growing, we arrested the simulation at different computational time points (Figure S1). We observed a similar trade-off among capacity, security and decoding reliability as mentioned in the main text (Figure S12). The patterns arrested at earlier time points have higher encoding capacity and lower security, as they require fewer training data to achieve the same accuracy compared to the later patterns. This is possibly because as the patterning process goes on, the colony starts developing branches that obscure the initial configuration information and shrinks the inter-categorical similarity. The accuracy curves are bounded from below by the curve of the final patterns, implying that the final patterns provide the highest security among all. Moreover, the results indicate that temporal regulation of pattern formation would be an additional strategy to modulate encoding capacity and security.

**Authenticating patterns using noise signatures**

Information integrity is critical for reliable communication and could be compromised under different scenarios. For instance, the attackers could alter the patterns to prevent delivering important messages or replace them with fake ones to deceive the intended recipient (ex. phishing). Therefore, it is critical to carry out an integrity check before decoding. One plausible method is through hashing, where a unique hash is generated and assigned to a message. When the pattern is tampered, the hash should change drastically (so-called *avalanche effect*) and fail to match the original one, indicating potential attacks and damage. Moreover, the hash function should be designed so that the chance of two different patterns having the same hash value (i.e., collision) is low.

In our system, branching dynamic amplifies the inherent biological noise in cell seeding and growth, resulting in patterns that are similar globally but vary in detail. We leveraged this feature and implemented hashing in the pattern-based communication platform. For a given pattern, we first locate its bifurcation and branch ridges (Figure S13A). To compute a unique hash, we input the extracted minutiae locations into a symmetric hash function(2). This class of hash functions has the advantage of being order-invariant, such that it allows us to bypass the challenge of assigning orders to minutiae. Consider an extracted pattern minutia has location $\left( x_{i}, y_{i} \right), i\in\left\{ 1,2, \ldots, N \right\}$, where N is the total number of minutiae. We can design hash function $h_{k}=\sum_{i=1}^{N} (x_{i}^{k}+y_{i}^{k})$, where k indicates the order of the function. As a proof of principle, we tested hash functions of different orders on an example dataset and found that it was possible to find order k that minimizes the chance of collision for patterns encoding different characters (Figure S13B) and patterns encoding the same characters (Figure S13C). Moreover, it is also possible and probably more robust to match the patterns by comparing multiple hash values all at once.

In practice, the end-users can use the hash values to carry out pattern integrity checks before preceding to decode, similar to two-factor authentication (Figure S13D). Here, we focused only on the malicious tampering of patterns other than transformation or numerical errors caused by minor transmission distortion. Considering the similarity between matching branching patterns and human fingerprints, further studies could design and apply more robust hash functions, such as those used on biometric data(3-5).

**Encoding and decoding using Elementary cellular automaton**

The elementary cellular automaton (ECA) model is one-dimensional, each cell has a state of either 0 or 1. The system starts from an initial sequence of cells, defined in a similar manner as in the Deng model (Figure S4C). Each character is first converted into a binary number, which is then translated into an initial configuration according to a predefined array. The cell status at the next time step depends on the current status of its neighbors. We implemented rule 60 that is weakly chaotic. Here 60 represents binary number 00111100, each digit represents the resulting cell status in the corresponding scenario (Figure S4A). For example, in the first scenario (Figure S4A, the most left), the cell of interest (middle) and its two neighbors have status of 111 at the current time step, the first 0 in the binary number means the cell of interest will have status 0 at the next time step.

Unless specified, the default encoding parameters are: sequence length = 450, time step = 500, spot radius = 10, dictionary size = 15, spacing = 70. Noise is imposed onto the seeding sequence before evolution starts. We draw random numbers from a uniform distribution and assign them to each cell within the initial configuration. We define a threshold ρ, cells with value smaller than ρ will flip from status 1 to 0, otherwise they will keep status 1. In other words, ρ is the percentage of cells who flip their status, larger ρ indicates larger noise level. We use default ρ of 0.5. These parameters were chosen such that the system gives satisfying security level. To decode, we trained a feedforward neural network to classify the output sequences via classification. The mathematical modeling was implemented in MATLAB R2020b, the ML model training implemented in Python3 and PyTorch 1.9.

Figure S5 shows that increasing noise level and small spacing deteriorate decoder performance, which is consistent with the observations with colony patterns. The decoding accuracy can be partially rescued by increasing the training data size. Dictionary size does not impact the performance, indicating each system have different encoding capacity and must be tuned individually.

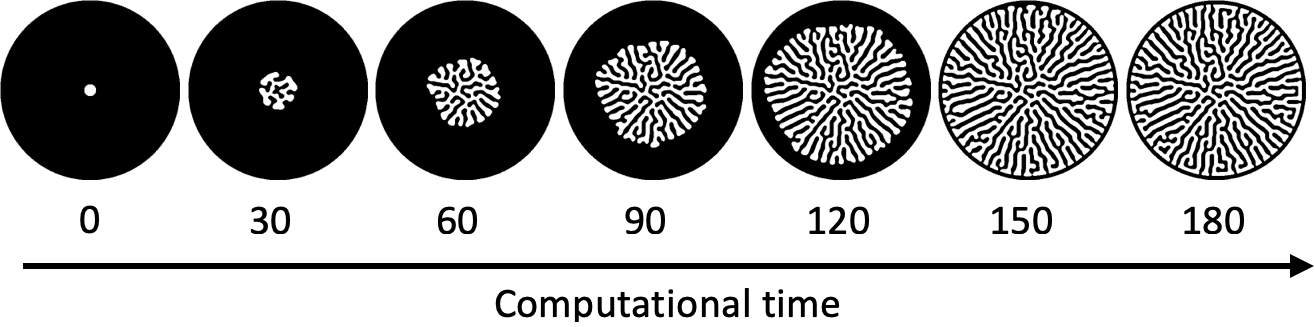

**Fig. S1.** Time series of branching pattern development.

An example of colony growth starting from a single spot configuration at the center of a circular growth domain. The cells are unevenly seeded within the initial configuration. As the colony grows, it develops into an intricate branching pattern and stops growing at around t = 180. The simulation parameters are the default: seeding spot radius = 5, $d_{1}/d_{2}=0.4 , h_{1}=1000, h_{2}= 2000, b=6.5$, $\epsilon=2000$, there was not growth noise (See “Methods”).

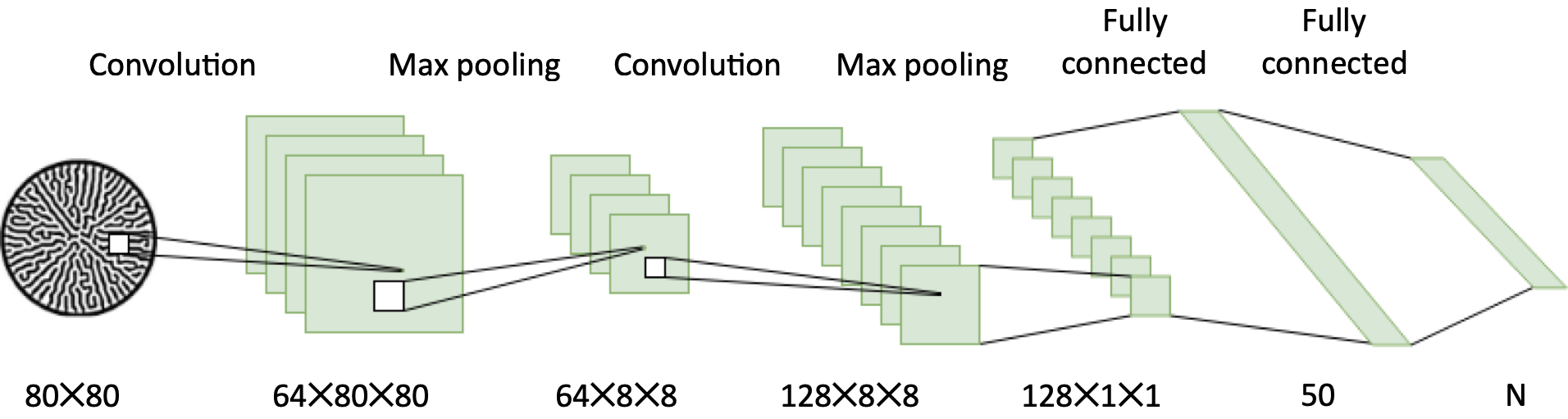

**Fig. S2.** Schematic of CNN decoder architecture.

The CNN takes an 80 × 80 greyscale image as input, and outputs a N dimensional feature, where N is the number of characters in a dictionary.

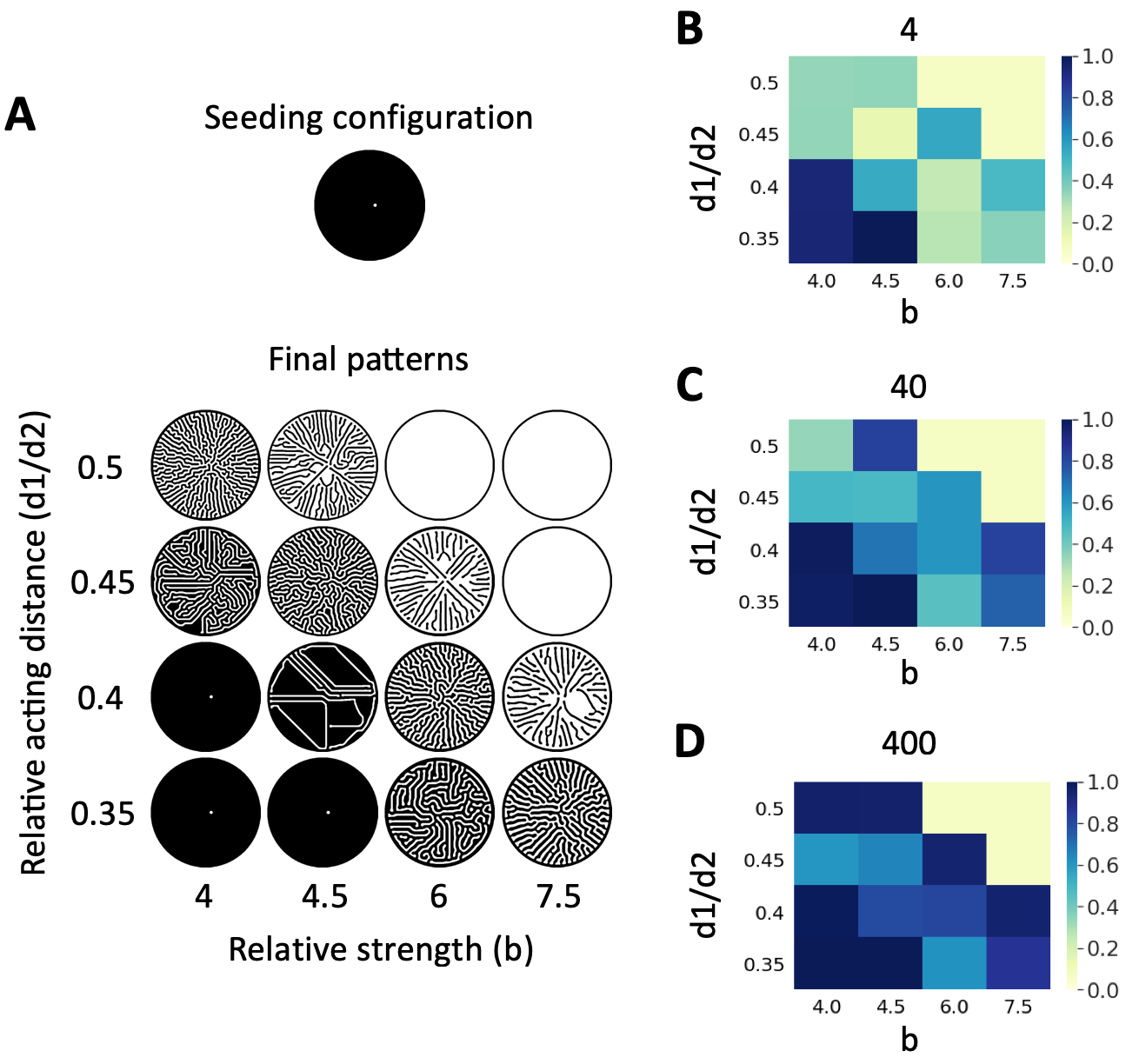

**Fig. S3.** Patterning dynamics impact the tradeoff among encoding capacity, security, and decoding reliability.

A. Starting from the same initial seeding configuration (and the same seeding noise), cells grow into diverse final patterns under different growth dynamics. Here, the growth dynamic is defined by two parameters: the relative strength and the relative acting distance of colony expansion and repulsion. The simulation is terminated when the growth stops. We can coarsely classify the patterns into three groups: the trivial (lower left corner), the disk-like (top right corner) and the branching patterns (the ones along the diagonal).

B – D. Decoding accuracy of different dynamics when scarce (B), intermediate (C), and adequate (D) training data are available (4, 40, and 400 replicates per class respectively). The trivial patterns always have high decoding capacity, whereas the disk-like patterns always have low decoding capacity. For branching patterns, the decoding accuracy increases drastically if more data become available.

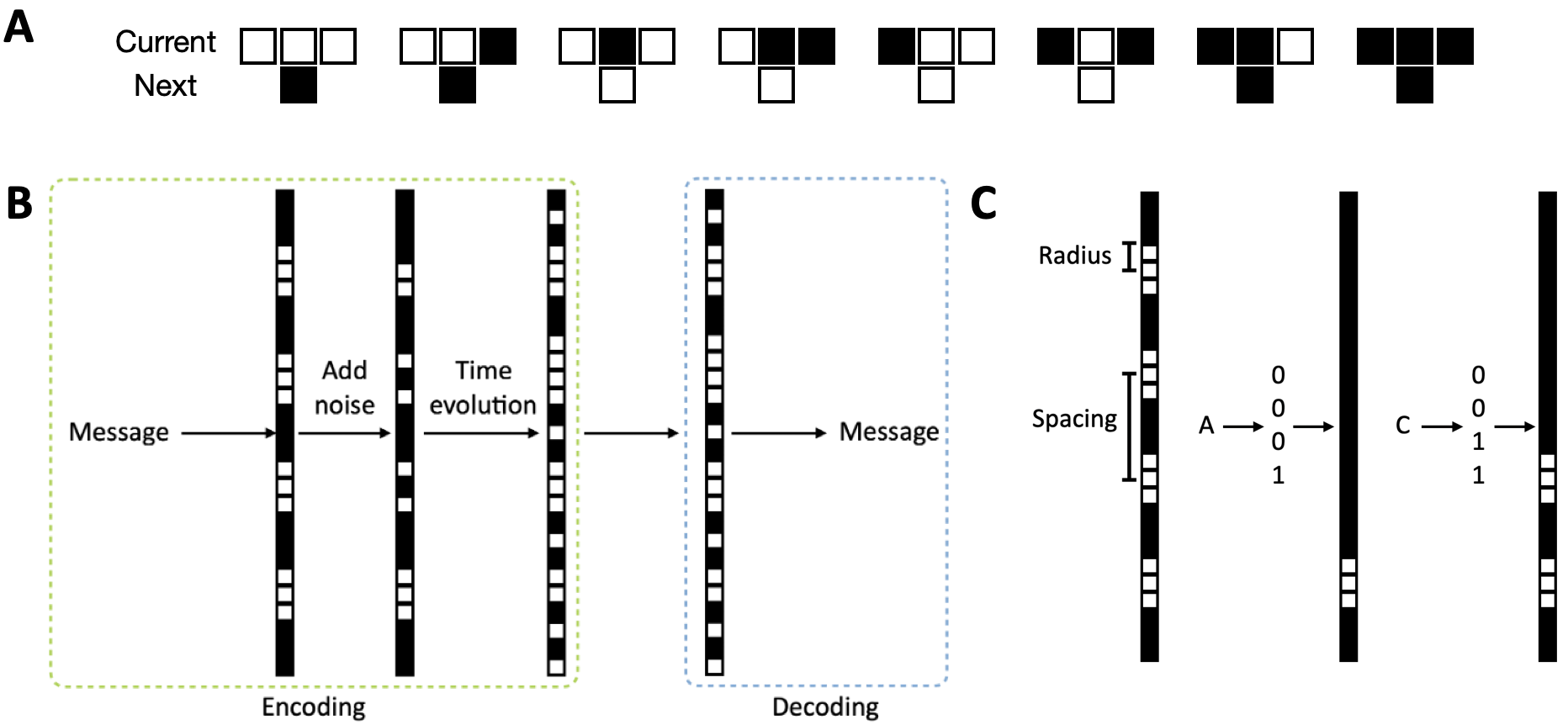

**Fig. S4.** Encoding and decoding using elementary cellular automata (ECA) with rule 60.

A. Rule 60 of ECA. Color indicates cell status: black – 0; white – 1. For the specific “Current” states, the sequence of the "Next” states represents the number 60 in a binary format (00111100).

B. Encoding-decoding scheme using one dimensional ECA. To encode, a message is first converted into a one-dimensional seeding array and then noise is added to it. The sequence then evolves into a final pattern following rule 60. A trained feedforward neural network is used to decode the pattern.

C. Predefined braille-like cell seeding arrangement and examples of encoded letters.

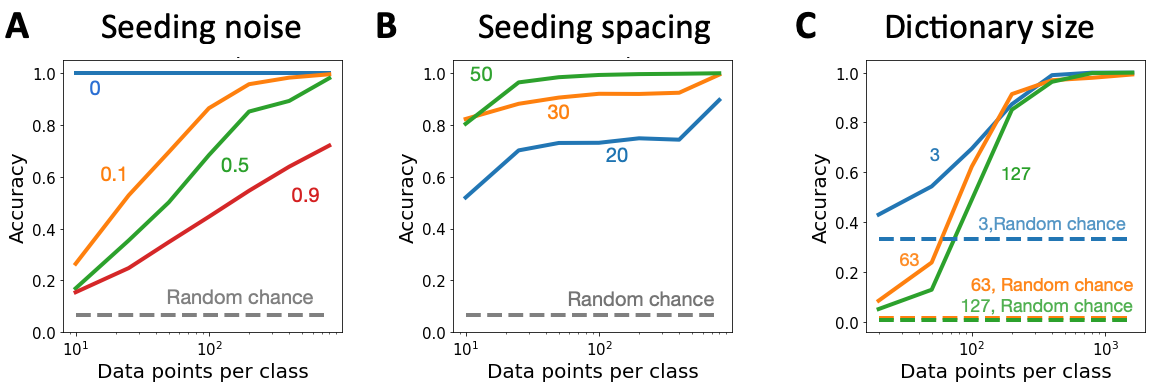

**Fig. S5.** Tradeoff between encoding capacity, security, and decoding reliability of the CA model. We varied parameters including: A. level of noise in time evolution (increasing value indicates higher noise), B. spacing between spots (time step = 600), and C. number of characters in encoding setup. The decoding accuracy generally increases as the number of replicates per class increases, and it significantly exceeds the corresponding accuracy by random guessing. Increasing complexity such as using larger growth noise or smaller spacing would require more data to reach the same accuracy. Larger dictionary size does not lead to sufficiently distinguishable training performance profile, it indicates every system may have different capacity and require tuning.

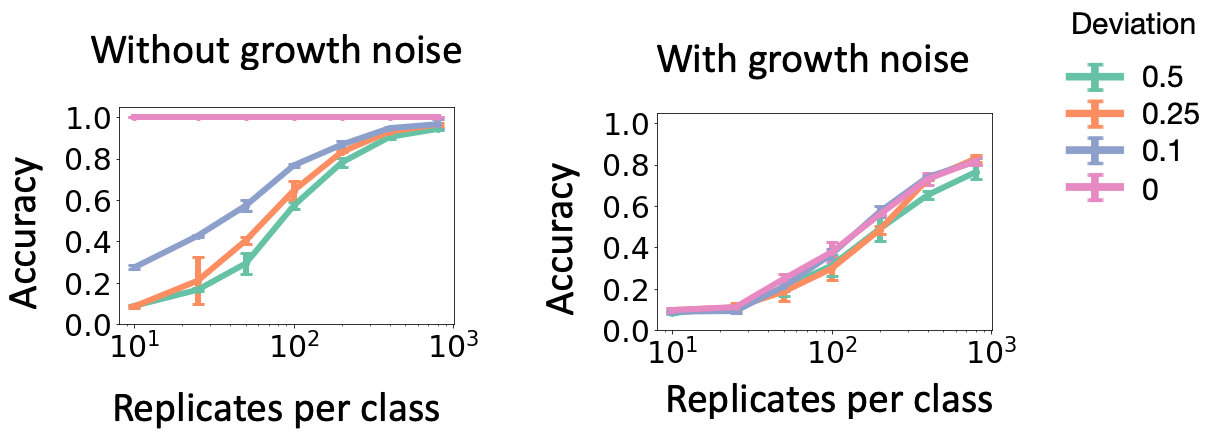

**Fig. S6.** Impact of seeding noise on the tradeoff among encoding capacity, security, and decoding reliability.

The seeding noise is implemented by assigning each pixel within the seeding configuration a random value. These values are drawn from a truncated Gaussian distribution with a mean of 0.5 and a given deviation (0, 0.1, 0.25 and 0.5 respectively). Larger deviation results in larger seeding noise. In the absence of growth noise, larger seeding noise leads to more challenging decoding, which is indicated by the increasing required training data. However, this impact is minor and can be obscured by a gentle growth noise (SNR = 10, see “Methods” for more details).

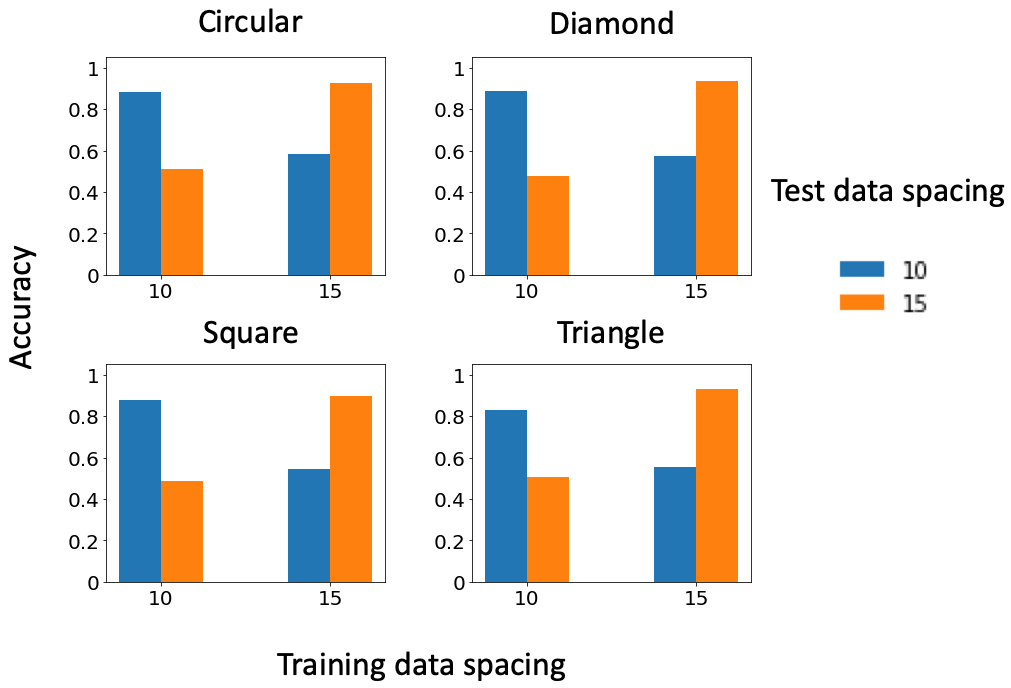

**Fig. S7.** Spacing as the encryption secret key.

Spacing distance = 10 and 15 were used respectively as the secret key to generate the training patterns. On growing media of all shapes, only models trained with the correct dataset can decode the patterns at significantly higher accuracy. The results indicate that spacing is a feasible choice for the secret key.

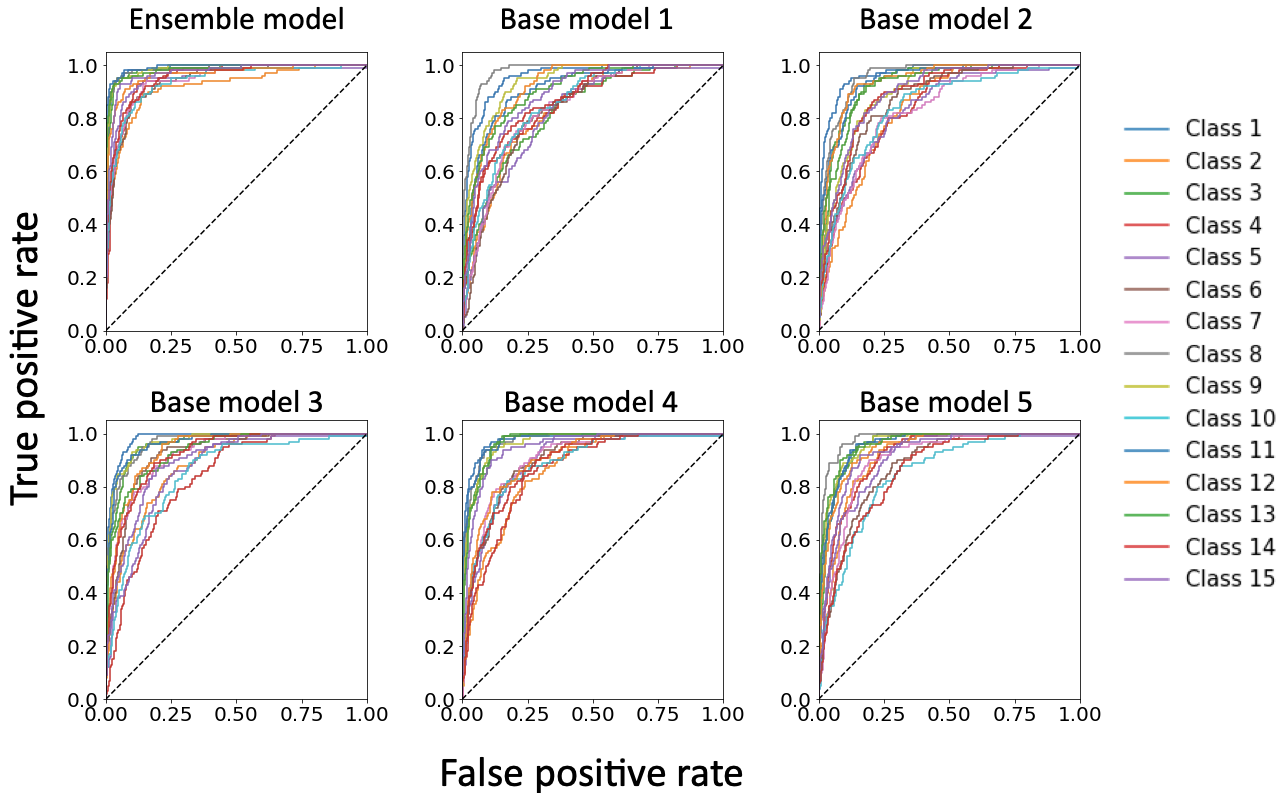

**Fig. S8.** Multiclass ROC of ensemble and base models.

We used 5 base CNN models to train a LR ensemble model. The training data were patterns generated using moderate growth noise (SNR = 3.5) and the dataset was composed of 100 replicates per class. Among base models, the ROC curves vary drastically. On the contrary, the variation reduces for the ensemble model, and the curves shift towards the upper left corner indicating significant performance improvement.

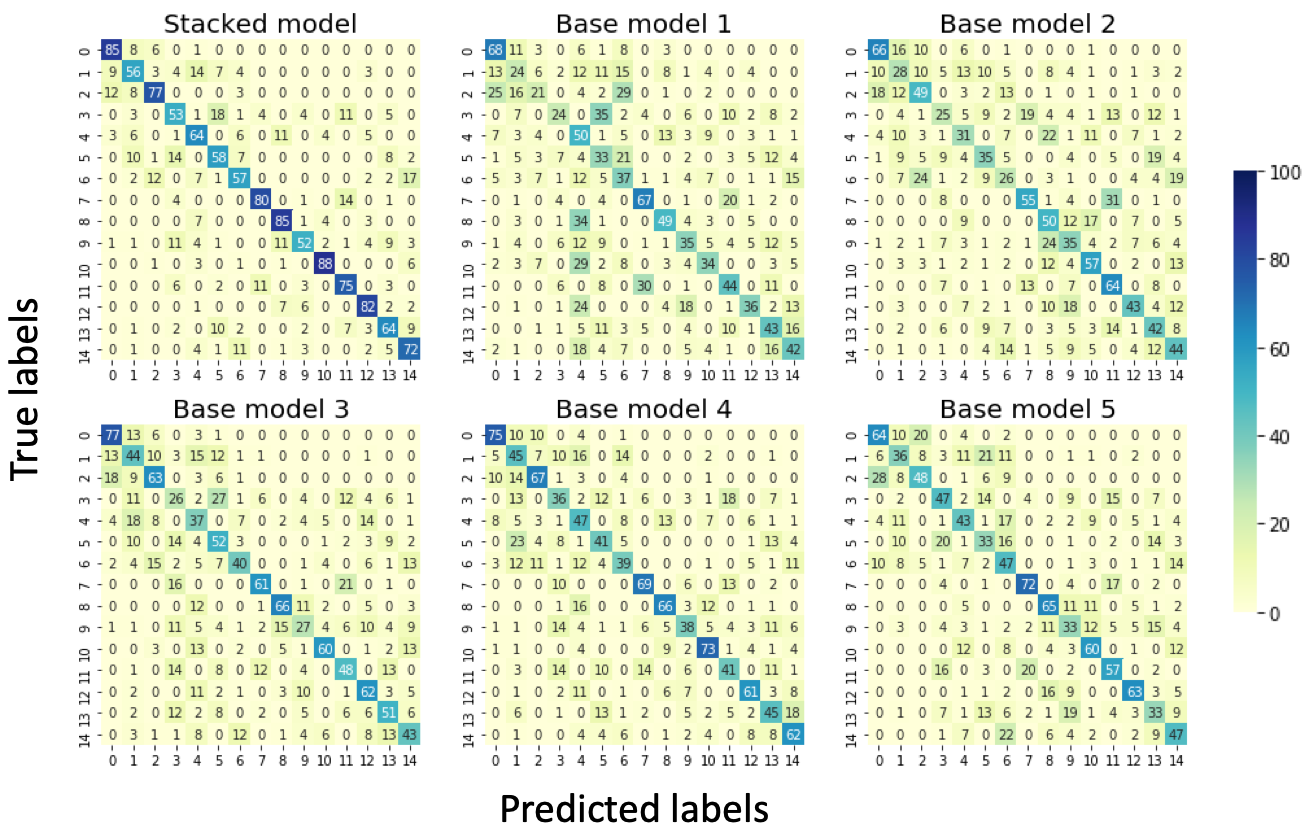

**Fig. S9.** Confusion matrix of ensemble and base models.

The base and ensemble models were trained on the same dataset in the same way as in Figure S6. Overall, the chance of misclassification reduces considerably for the ensemble model in comparison to individual base models.

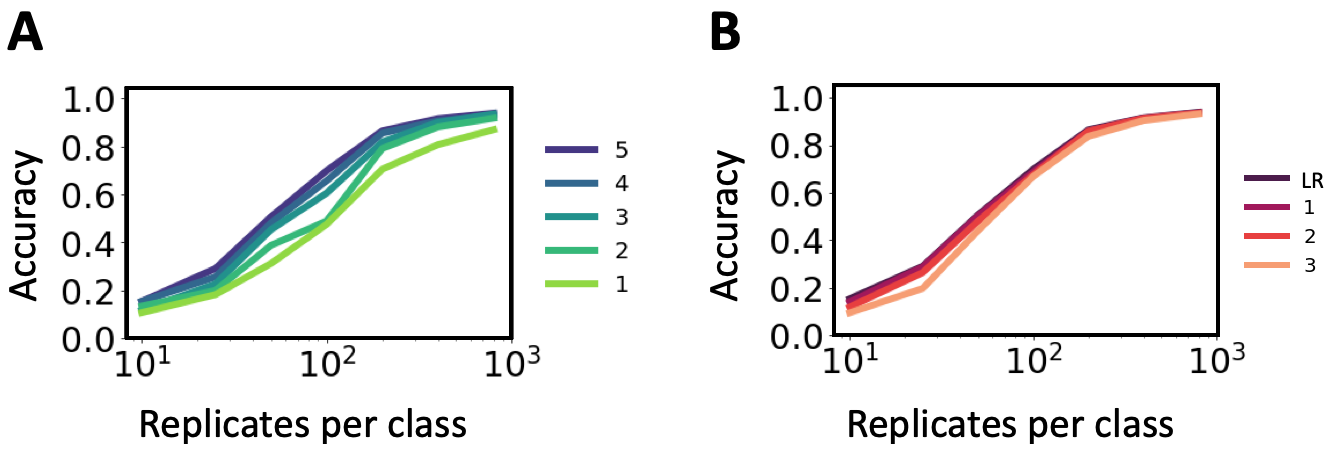

**Fig. S10.** The impact of the number of base models and ensemble model architecture on ensemble prediction.

A. Colors indicate different number of base models. The ensemble accuracy improves as more base models are used for constructing a LR ensemble model, and the amount of increase depends on the dataset size. The accuracy would eventually saturate when sufficiently many base models are used.
B. Logistic regression (LR) as the ensemble model outperforms feedforward neural network (FNN) with either 1, 2 or 3 hidden layers. Here, we kept the input and output layers the same for all models, and the number of hidden nodes for the FNNs are (40), (60, 30) and (60, 45, 25) respectively. LR performs slightly better or as good as FNNs.

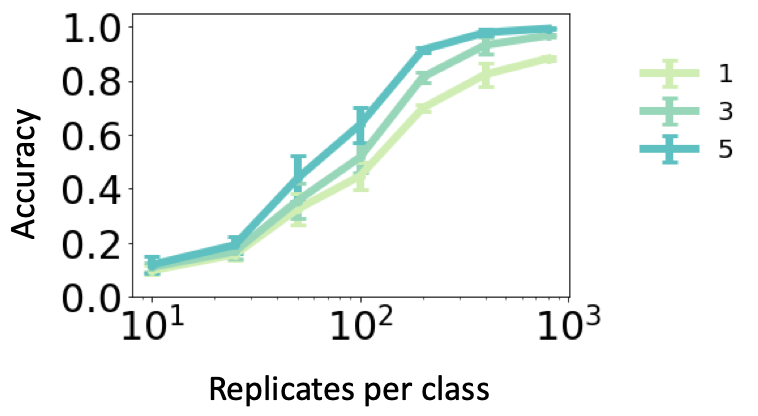

**Fig. S11.** Majority voting improves decoding accuracy.

Decoding accuracy of using 1, 3 and 5 patterns under the majority voting scheme. Using more patterns can significantly improve the performance, especially when a large dataset has been used for training.

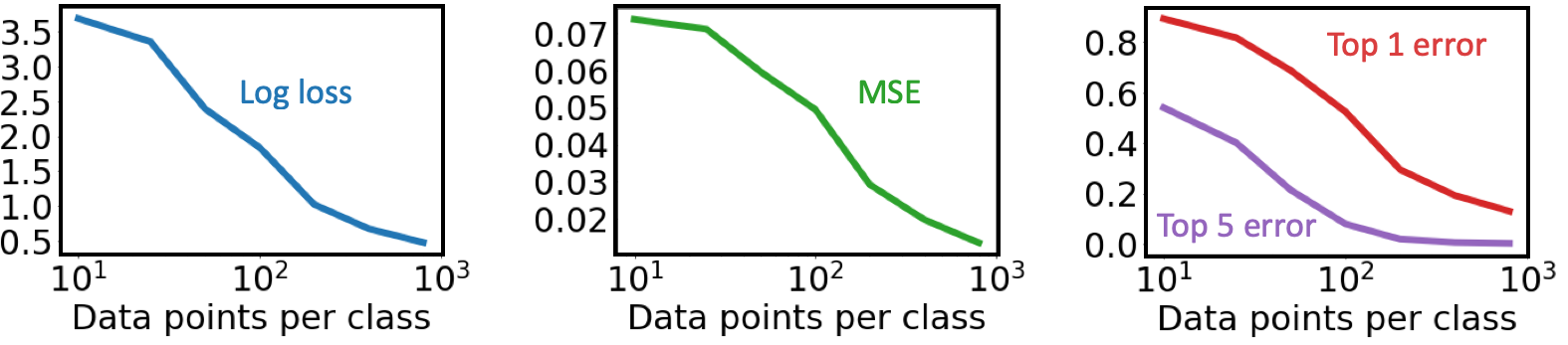

**Fig. S12.** Uncertainty estimation using deep ensemble.

5 base models trained using random initialization were used to calculate the selected metrics. Higher value indicates greater uncertainty. Having more training data lowers the prediction uncertainty.

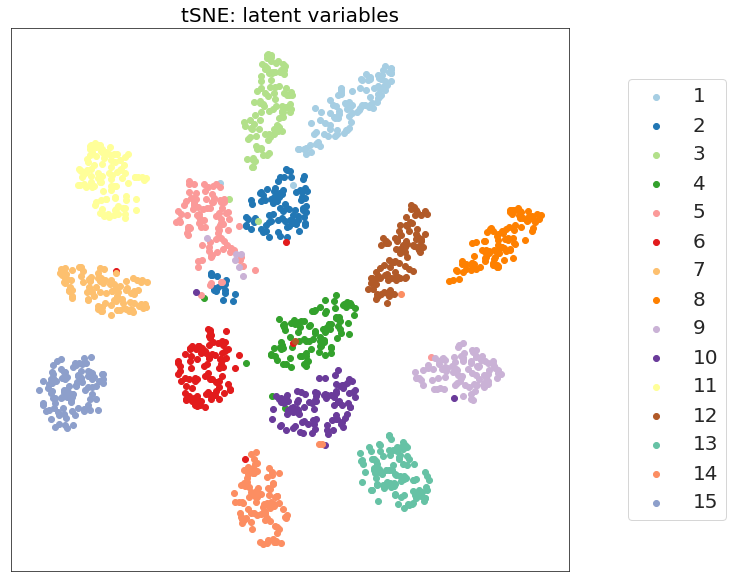

**Fig. S13.** t-distributed Stochastic Neighborhood Embedding (t-SNE) of data distribution.

t-SNE was trained to embed CNN outputs (before fed to softmax) into 2D space using random initialization, perplexity 50, and learning rate 200 (implemented in scikit-learn). The training dataset consists of 15 initial configurations and 12000 data points, the simulation parameters are the default. The figure illustrates the distribution of 100 randomly selected patterns from each initial configuration (each labeled by a color). The results show that the patterns encoding different characters are indeed distinguishable as t-SNE learnt to cluster them.

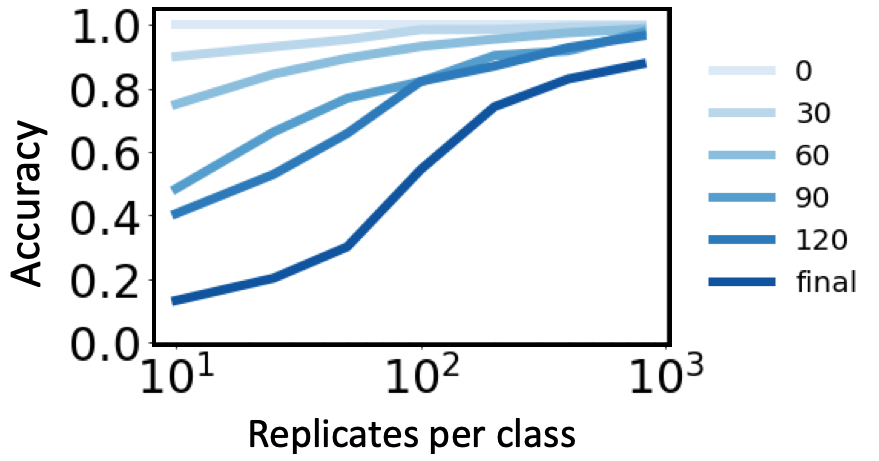

**Fig. S14.** Temporal information encoding using branching patterns.

Decoding accuracy on patterns arrested from growth at t = 0, 30, 60, 90, 120 and when the colony stops expanding (“final”). The patterns stopped growing at earlier time points, require less training data to achieve the same decoding accuracy.

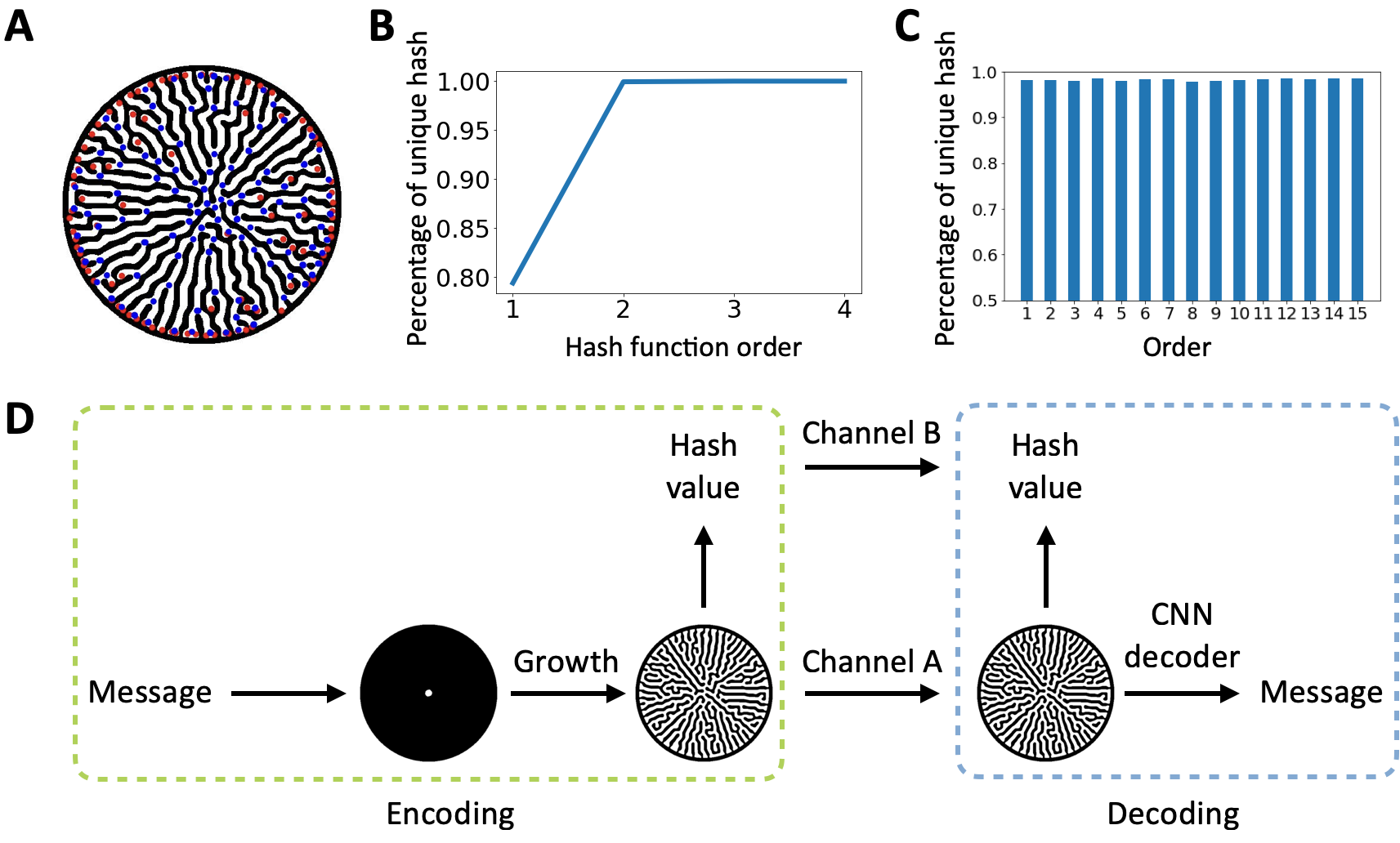

**Fig. S15.** Protecting information integrity using biological noise

A. Extracted minutiae (blue: bifurcation, red: branch ridge) of an example pattern.
B. The percentage of unique hashes. The hashes were computed using symmetric hash functions with order k = 1, 2, 3, and 4, respectively on a dataset of 15000 patterns (15 classes, 1000 replicates of each class). The default parameter set was used for data generation. To compute the percentage, we removed 217 duplicate patterns and normalized the counts with respect to the total number of unique patterns (14783). The figure indicates that increasing the order of hash function can reduce the chance of two different patterns sharing the same hash.
C. The percentage of unique hashes by class. The hash function here has k = 1. The calculation was carried out on the same dataset as in panel B. The percentage of unique hash was normalized with respect to the number of unique patterns for a given class. The results show that the chance of collision is comparable among all classes.
D. Integrity check procedure. In real communication, the sender would send a pattern and its hash through different channels (channel A and B, respectively) to the recipient. Once the recipient receives both, he/she would first authenticate information integrity by computing the hash of the received pattern and comparing it with the hash received through channel B. If they agree, the recipient can proceed to decode. Otherwise, reject the pattern and request a new one. This procedure is analogous to a typical two factor authentication.

**Table S1.** Converting encoding characters into binary representations.

For an example dictionary of 15 characters (A-E, 0-9), one possible way of encoding is to first convert them into 4-bit binary numbers in order. Binary number “0000” is purposefully avoided, because 0 corresponds to the absence of cells in the seeding configuration, thus colony patterns cannot form.

| Encoding character | Binary representation | Encode character | Binary representation |
| --- | --- | --- | --- |
| A | 0001 | 3 | 1001 |
| B | 0010 | 4 | 1010 |
| C | 0011 | 5 | 1011 |
| D | 0100 | 6 | 1100 |
| E | 0101 | 7 | 1101 |
| 0 | 0110 | 8 | 1110 |
| 1 | 0111 | 9 | 1111 |
| 2 | 1000 |  |  |

**Table S2.** Estimated uncertainty as a function of training data size and the number of base models.

We trained LR ensemble model using differently sized dataset and number of base models. For the same dataset, the estimated uncertainty does not change drastically with respect to the number of base models.

| Replicates per class | Number of base models | LL | MSE | Top 1 error | Top 5 error |
| --- | --- | --- | --- | --- | --- |
| 10 | 5 | 3.687 | 0.074 | 0.107 | 0.541 |
|  | 4 | 3.675 | 0.074 | 0.108 | 0.537 |
|  | 3 | 3.645 | 0.073 | 0.098 | 0.552 |
|  | 2 | 3.545 | 0.072 | 0.108 | 0.533 |
| 100 | 5 | 1.839 | 0.049 | 0.476 | 0.080 |
|  | 4 | 1.854 | 0.050 | 0.470 | 0.088 |
|  | 3 | 1.903 | 0.051 | 0.448 | 0.097 |
|  | 2 | 2.010 | 0.054 | 0.419 | 0.110 |
| 800 | 5 | 0.478 | 0.014 | 0.869 | 0.004 |
|  | 4 | 0.423 | 0.012 | 0.881 | 0.003 |
|  | 3 | 0.425 | 0.012 | 0.882 | 0.003 |
|  | 2 | 0.446 | 0.012 | 0.884 | 0.002 |

**Movie S1 (separate file).** Video encoding “I have a dream”.

A 7-bit spot seeding array was used to generate a dataset containing 127 classes and 1000 replicates each. Each character was represented by a randomly chosen image from the dataset, arranged in the same order as the text. We repeated this process 5 times and attached each random array of images after another. To decode, we trained a neural decoder on the dataset, extracted every frame, and decoded them one by one. The 5 patterns representing the same character were used in majority voting, the results of which was used as the final prediction. Due to file size limitation, this video only contains part of the entire speech (“Five score years ago, … every American was to fall heir.”) using compressed 80 × 80-pixel pattern images.

**Movie S2 (separate file).** Video encoding “Auguries of Innocence”.

The video represents the poem “Auguries of Innocence” by William Blake. Due to file size limitation, this video only contains part of the entire poem (“To see a world … in endless Night”) using compressed 80 × 80-pixel pattern images. The encoding and decoding were conducted using the same protocol as for Movie S1.

**Movie S3 (separate file).** Video encoding GFP sequence.

A 5-bit spot seeding array was used to generate a dataset containing 31 classes and 1000 replicates each. Among the 31 classes, 20 of them were used to represent the common amino acids. The encoding and decoding were conducted using the same protocol as for Movie S1. The video represents the complete GFP sequence and is consisted of compressed 80 × 80-pixel pattern images.
